# Revisiting biogeography of livestock animal domestication

**DOI:** 10.1101/786442

**Authors:** Indrė Žliobaitė

## Abstract

Human society relies on four main livestock animals – sheep, goat, pig and cattle, which all were domesticated at nearly the same time and place. Many arguments have been put forward to explain why these animals, place and time were suitable for domestication, but the question – why only these, but not other animals, still does not have a clear answer. Here we offer a biogeographical perspective: we survey global occurrence of large mammalian herbivore genera around 15 000 – 5 000 years before present and compile a dataset characterising their ecology, habitats and dental traits. Using predictive modelling we extract patterns from this data to highlight ecological differences between domesticated and nondomesticated genera. The most suitable for domestication appear to be generalists adapted to persistence in marginal environments of low productivity, largely corresponding to cold semi-arid climate zones. Our biogeographic analysis shows that even though domestication rates varied across continents, potentially suitable candidate animals were rather uniformly distributed across continents. We interpret this pattern as a result of an interface between cold semi-arid and hot semi-arid climatic zones. We argue that hot Semi-arid climate was most suitable for plant domestication, cold Semi-arid climate selected for animals most suitable for domestication as livestock. We propose that the rates of domestication across biogeographic realms largely reflect how much intersection between hot and cold Semi-arid climatic zones was available at each continent.

## 1. INTRODUCTION

One of the fundamental questions in human history is why so relatively few animal and plant groups have been domesticated (Diamond 1997). Human society today relies on four main livestock animals (sheep, goat, pig and cattle), which all were domesticated at nearly the same time and place – starting about 10 500 years ago at the Fertile Crescent^1^. Even more intriguing is that those species did not evolve locally at or near the Fertile Crescent, the wild progenitors immigrated mostly from the Central Asia (Diamond 1997). Curiously, domestication of those livestock animals happened repeatedly at the same place, as “genetic analyses report multiple domestic lineages for each species” (Zeder 2008). Many arguments have been put forward to explain why these animals, place and time were suitable for domestication (see e.g. Diamond 1997, Larson and Fuller 2014, Zeder 2017, MacHugh et al 2017, Crosby 2006). But key questions – why animals from elsewhere were more suitable for domestication instead of native animals, why those animals have not been domesticated at their places of origin, and why animals local to the Fertile Crescent have not been domesticated instead or in addition, largely remain unanswered.

The main objective of this study is to investigate to what extent opportunities for early livestock domestication have been exhausted. To analyse options for animal domestication, we ask what potential candidate species were available around the time of first known domestications of livestock, given in Table 1. We analyse ecological, biogeographic and climatic context of large plant eating mammals contemporary to the time of those early domestications aiming to deduce common patterns of their physiology, ecology, dietary preferences and inferred behaviour. We computationally model the probability of domestication as a function of those factors. To the best of our knowledge this is the first quantitative analysis of domestication patterns at the global scale. We make the analysis datasets publicly available for research and development.

**Table 1.** Domestication of livestock animals. Domestication time is an approximate time of transition between management of animals and morphological changes associated with domestication, rounded to the nearest thousand. The list of animals is derived from Diamond (1997), Larson et al (2014), and Zeder (2008).

| Domestic animal | Ancestor | Domestication place | Time, B.P. | Digestive type |
| --- | --- | --- | --- | --- |
| Sheep | Asiatic mouflon | The Fertile Crescent | 10 000 | Ruminant |
| Goat | Bezoar ibex | The Fertile Crescent | 10 000 | Ruminant |
| Pig | Eurasian wild boar | The Fertile Crescent | 10 000 | Non-ruminant |
|  |  | China | 8 000 |  |
| Cattle (taurine) | Aurochs | The Fertile Crescent | 10 000 | Ruminant |
| (zebu) |  | South Asia | 8 000 |  |
| Horse |  | Central Asia | 6 000 | Non-ruminant |
| Donkey | *Equus africanus* | North Africa | 5 000 | Non-ruminant |
| Llama |  | South America | 5 000 | Pseudoruminant |
| Alpaca | *Vicugna* | South America | 4 000 | Pseudoruminant |
| Water buffalo | *Bubalus arnee* | South Asia | 4 000 | Ruminant |
| Yak | *Bos mutus* | Central Asia | 4 000 | Ruminant |
| Camel (Bactrian) | *Camelus ferus* | Central Asia | 4 000 | Pseudoruminant |
| Camel (Dromedary) |  | North Africa | 3 000 | Pseudoruminant |

<sup>1</sup> The Fertile Crescent is a region in the Middle East, that lays across the territories of present-day Iraq, Israel, Syria, Lebanon, Egypt and Jordan, it reaches some parts of Turkey and Iran, and Cyprus may or may not be included. The name was popularized by James H. Breasted (1914).

## 2. MATERIALS AND METHODS

Our study region is the world at times of early livestock animal domestication, we consider a time range from around 15 000 to 5 000 years before present. Domestication has many definitions and interpretations (Decory 2019) that vary across groups of organisms and contexts of usage. Here we consider domesticated those species for which humans have significant degree of influence over the reproduction and care, and which have undergone significant genetic, behavioural and/or morphological changes since. Here we consider only livestock animals and do not cover birds.

Our study consists of two parts: analysis of ecological characteristics of candidate animals, and biogeographic context of their domestication. For the latter, we divide the world into six regions following designation of biogeographic realms at the present day (Olson et al 2001): Nearctic (mainly North America), Palearctic (mainly Eurasia), Afrotropic (mainly Africa), Indomalaya (South East Asia), Australasia and Neotropic (South and Central Americas). Mapping of those realms is given in the Supplement. The realms distinguish large terrestrial areas within which organisms have been evolving in relative isolation over long periods of time, separated by geographic features, such as oceans, broad deserts, or high mountain ranges. We therefore use realms as natural units for analysing and comparing domestication rates across the globe.

### 2.1. The dataset of candidate genera

We compiled the list of candidate genera for domestication following the lists of living mammalian species (Wilson and Reeder, 2005, and Nowak, 2018), assuming that genera that are alive today were present in the same biogeographic realms within the selected time frame in the past. We complemented our list of candidates by fossil genera from our own archives and the NOW database of fossil mammals (The NOW Community, 2018). We excluded genera that went extinct before 5000 B.P., assuming that they were rare at the times of possible first domestications (e.g. *Hippidion* in South America). Even though taxa could have been domesticated just before they went extinct, while going towards extinction rarity of individuals increases (Zliobaite et al 2017) and perhaps persistence in native environments is becoming more challenging, even if domestication is attempted. Our analysis of is at the level of genera rather than species, since genera is a more robust unit of analysis for the past (Eronen et al 2011) and cogeneric species rarely co-occur anyway (Levin et al 2012).

We chose to compile a list from scratch rather than building on existing regional list aiming for consistency of treatment and due to lack of existing options. A regional compilation of candidate species for South-West Asia by Garrad (1984) exists, which would have covered only a small part of our scope. Diamond (1997) compiled a global list, which he used for computing domestication rates at continents, but the list of candidate taxa was not made available in his publication.

Our candidate list included herbivorous genera within the body mass range from 40 to 1000 kg, which covers the body mass classes of early domesticated livestock animals. Having livestock animals large enough saves the need to herd countless numbers of individuals. Yet domestic animals cannot be too large, since otherwise it would be too challenging to handle them and it would take too much time and effort to raise them to maturity. Therefore, neither small mammals, nor megaherbivores, such as elephants, are particularly practical for domestication. We did not include carnivores, but included omnivores. In retrospect, the most suitable seem to be large herbivores, from about human size to at most an order of magnitude larger. This goes in line with carnivore-prey body size relationships in mammals, where preferred prey is from about half to about twice size of the carnivore (Owen-Smith and Mills 2008). We assume that the lower bound for humans is the same as for predators hunting in the wild, but the upper bound for humans can go higher because of tools and technology available for humans for managing and killing livestock animals.

We characterized each candidate taxa by their habitat, dietary characteristics, and functional dental traits, which made 15 features in total aiming to capture their ways of life. Habitat and dietary characteristics were collected from numerous academic sources and animal databases. We did not record individual sources for the habitat and dietary variables, and cannot provide referencing for each individual data point thereof. Given our choice to work at the resolution of genera, habitat and dietary variables can only be approximate, since a genus may include species that have different diets or habitats, or their ways of life may have changed between 15 000 years before present and now. We resolved within genus variations by assigning characteristics of the most common species, or, in rare cases of equal balance of the opposite characteristics, we assigned a half value. If information about the behaviour now and in the past diverged, we assigned the value for the past. We characterized dental traits following the functional crown type scoring scheme described by Zliobaite et al. (2016) with one modification allowing selenodonts have acute lophs, following the reasoning of Oksanen et al (2019).

Each candidate genus was described by 15 binary or ordinal variables, given in Table 2. All the variables except for hypsodonty were encoded in the scale from 0 to 1. We kept hypsodonty in the original ordinated scale (Fortelius et al., 2002), in order not to introduce extra complications in interpretations. A detailed description of the scoring scheme is given in the Supplement. A full dataset along with the list of genera excluded from the candidate list and reasons thereof are given in the Supplement as well.

**Table 2.** Variables describing characteristics of the candidate genera.

| Variable | Possible values |
| --- | --- |
| 1. Grazer | 1 grazer, 0.67 grass-dominated mixed-feeder, 0.33 mixed-feeder that takes some grass, 0 no grass. |
| 2. Highly specialized | 1 highly specialized (e.g. aquatic) diet, 0.5 somewhat specialized, 0 common grazing or browsing. |
| 3. Water dependence | 1 drinking water daily, 0.5 every 2-3 days, 0 a week or more without water. |
| 4. Open habitat | 1 grassland, open savanna, steppe, 0.5 woody savanna, bushland, 0 dense forest. |
| 5. Highland habitat | 1 mountains or hills, 0.5 foothills, 0 steppes or forests. |
| 6. Migration | 1 regular seasonal migration, 0.5 seasonal change in range, 0 no migration. |
| 7. Highly endemic | 1 highly endemic, 0 widely distributed. |
| 8. Tropical | 1 tropical, 0 non-tropical (temperate or mountainous) habitats. |
| 9. Hypsodonty (HYP) | 3 hypsodont, 2 mesodont, 1 brachyodont. |
| 10. Acute lophs (AL) | 1 acute lophs present in molars, 0 absent. |
| 11. Obtuse lophs (OL) | 1 obtuse lophs present, 0 absent. |
| 12. Fortification (SF) | 1 structural fortification of molar cusps, 0 no fortification. |
| 13. Flatness (OT) | 1 occlusal topography flat, 0 non-flat. |
| 14. Goat-like (GO) | 1 obtuse lophs are present, but acute lophs, fortification or flatness are absent. |
| 15. Bunodont (BU) | 1 no lophs, 0 lophs of any type are present. |

As a summary of the dataset and a sanity-check, Figure 1 visualizes the first two (scaled) principal components of the dataset. We can see from the plot and the rotation matrix (given in the Supplement) that the primary axis is mostly hypsodonts (open habitats *or* grazing taxa) versus brachydonts (primarily forest dwellers). The second axis is primarily about arid environment vs. humid (such as swamps). Domesticated animals occur by and large in all spaces of this visualization, but the density of coverage is not uniform. Visually, the highest density of domesticated animals is at the arid (bottom-left) end, where camelids and goats are positioned.

**Figure 1.**
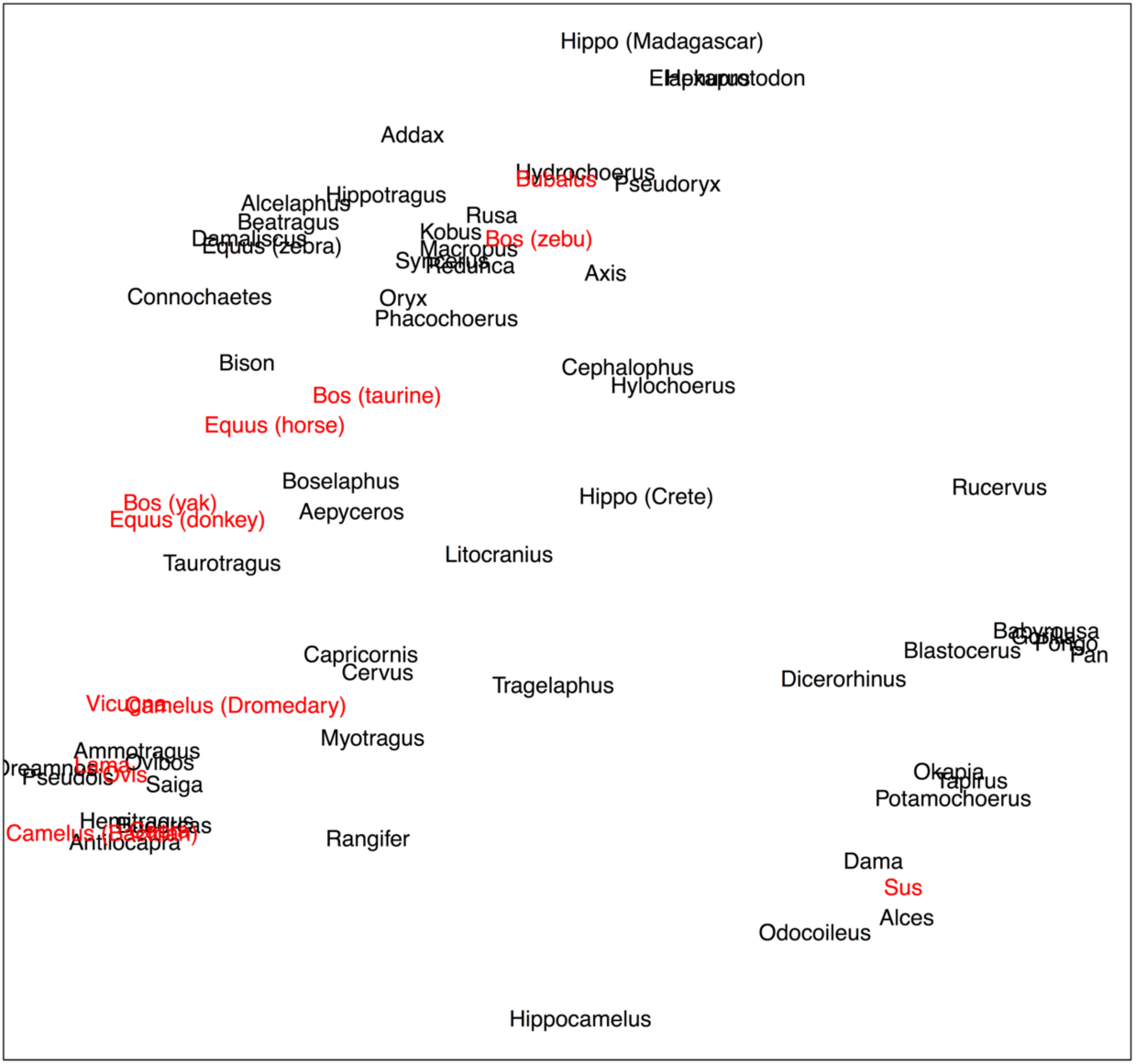
Visualization of two components with the highest variance resulting from singular value decomposition (of 15 input variables listed in Table 2). Domestication status is not included in the inputs for the projection.

### 2.3. Computational methods for predictive modelling

The main computational task is to learn predictive models for explaining patterns of domestication. In the machine learning terms, we need a probabilistic classifier of a model form where contributions of each variable would be easy to interpret. The inputs for the model can be any or all variables given in Table 2. The target variable is a binary class label indicating whether the taxon has been domesticated or not. One of the challenges for this predictive modelling is a relatively small sample size with a high imbalance of classes (13 domesticated taxa vs.55 not domesticated genera).

We used two types of models: logistic regression and decision tree (see e.g. Witten et al 2016 for an introductory text to machine learning). Logistic regression models a linear relationship between the input variables and the class label. The output is passed through an s-shaped function to make sure that the prediction falls between zero and one, and in our case, can be approximately interpreted as the probability of domestication. A number of alternative procedures for estimating the model parameters exist. We chose a combination of LASSO and Ridge optimization (alpha = 0.5, see e.g. Hastie et al 2009 for details). This selection keeps the regression weights from overgrowing and at the same time minimizes the number of variables selected to include into the final model. The number of variables included depends on internal statistical assessment of variance through the course of model fitting.

Decision tree is a non-linear model, which, conceptually similarly to LASSO, first selects the variable that explains the class the best, then second best and onwards. For fitting a tree, we used the standard CART algorithm with the Gini coefficient as the splitting criteria (see e.g. Witten et al 2011 for background technical information). We produced two decision tree models: one where all 15 variables are candidates, and the other without variables describing environment and habitat, the latter aimed to capture domestication patterns only from the organismal perspective. Our modelling choices are summarized in Table 3. Modelling was done in R software, using packages glmnet (for LASSO) and rpart (for tree fitting). The code used for the analysis is available on GitHub repository^2^, although not very well documented.

**Table 3.** Predictive models used for analysis and modelling choices.

| Model | Model fit | Variable selection | Parameter selection | Purpose |
| --- | --- | --- | --- | --- |
| Decision tree | CART, Gini split | Automated from all 15 variables | Cross-validation | Identifying factors that render livestock animals suitable for domestication |
| Decision tree | CART, Gini split | Automated from 10 variables | Cross-validation | Explaining domestication only by animals, no geography |
| Logistic regression | LASSO and Ridge | Automated from 14 variables | Cross-validation | Numeric estimates for suitability for domestication (ranking) |

<sup>2</sup> https://github.com/zliobaite/teeth-domestication

## 3. SUITABILITY FOR DOMESTICATION AND DOMESTICATION RATES

The purpose of our computational analysis is twofold: to identify which ecological and environmental characteristics best explain domestication of livestock animals, and to quantitatively describe each biogeographic realm in terms of estimated suitability of their candidate animals for livestock domestication 15 000 – 5 000 years ago.

### 3.1 Which ecological and environmental characteristics best explain domestication

Two inferred decision tree models are plotted in Figure 2. The first tree internally selected most informative variables from all 15 candidate variables in Table 2. The second tree was given only 10 candidate variables describing animals themselves (Grazer, Specialized, Water dependent and dental characteristics), excluding the variables describing environment or habitat (Open habitat, Highland habitat, Migration, Highly endemic, Tropical).

**Figure 2.**
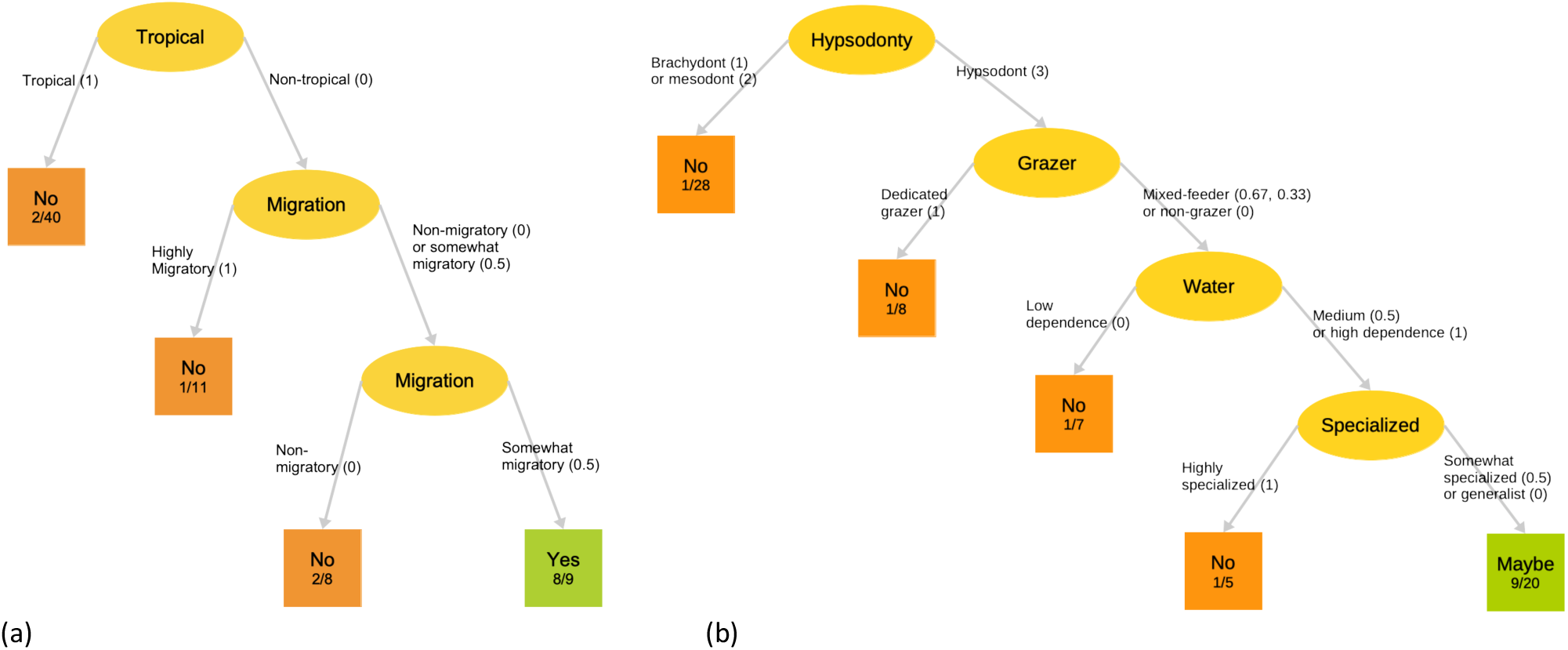
Decision tree models for domestication. (a) A model built using all the variables given in Table 2 as candidates. (b) A model built excluding environmental variables. The trees are read from top to the bottom: given a candidate animal, each yellow node indicates a variable to look at, and the values for a particular animal indicates which path to take until a terminal square node is reach, which gives an estimate of whether the animal is suitable for domestication. The number in the terminal nodes indicate how many animals with these characteristics have been domesticated and how many animals with these characteristics are in total in our dataset.]

The first tree can explain the majority of early livestock animal domestication cases (8 out of 13) by only two variables: coming not from tropical environments and being moderately migratory. This selection suggests that environmental criteria dominate over physiological in explanatory power. As the principle component analysis already hinted, domesticated animals appear through most of the space described by physiological traits, but the counts of successful domestication vary across that space. The highest densities of successful domestication, as the principal component and the tree analysis together suggest, must be the arid end of non-tropical environments. The fact that moderate migration has such a high explanatory power suggests a potential impact of cold and seasonality.

The second tree, which was not permitted to use environmental characteristics, gives a more entangled, but interesting pattern. Domesticable animals described by this tree are those having high hypsodonty, but not exclusively grazers. They are expected to be moderately or highly dependent on water sources, and not highly specialized (e.g. not aquatic animals). Yet, only half of the animals at the final node “Maybe” (9/20) were actually domesticated, which suggests that physiological characteristics alone, at least those included in our analysis, are not sufficient to explain the domestication patterns precisely. Realized domestication of livestock animals, seems clearly to be a combination of biogeographic and ecological factors.

Decision trees work well for visual and structural explanation, but without further modifications they are too coarse for probabilistic assessment of suitability of candidate animals for domestication. We need to assign suitability score for each candidate animal, such that we can rank them and reason about realized vs. potential domestication across biogeographic realms.

The inferred logistic regression model for estimating the probability of domestication is

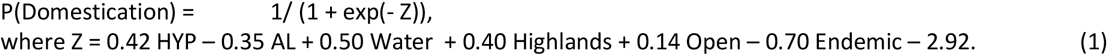

The model automatically selected the most informative variables from 14 candidate variables (all variables in Table 2 except for “tropical”, complementary model considering all 15 variables as candidates is given in Supplement). The regression component in the second line can be interpreted similarly as for a regular linear regression, where regression coefficients act as weights and their signs indicate the directionality of the relationship. The model predicts a high probability of domestication if an animal is hypsodont and does not have acute lophs, it is strongly associated with highlands and weakly with open environments, is not highly endemic, and it is somewhat dependent on availability of drinking water.

Judging from all three models the dominant, but not exclusive characteristics that make animals domesticable are their adaptedness to abrasive but not exclusively grass food (hypsodonty) in combination with somewhat marginal seasonally variable environmental conditions. This points to non-tropical generalists to have the highest propensity for livestock domestication.

### 3.2 Estimated suitability for domestication of candidate taxa

The next interesting question is how the 68 candidate taxa rank according to their potential suitability for domestication. Are there any taxa that within the context of other domesticated animals must have been very unlikely to be domesticated? And does any of the non-domesticated taxa look particularly suitable?

Table 4 gives a ranking of the candidates from the highest to the lowest probability of domestication. Probabilities are estimated using the logistic regression model given in Equation (1). For information, we also include scores given by both decision trees.

**Table 4.** Ranking of candidate taxa from the highest probability of domestication to the lowest estimated by the logistic regression model. For information probabilities estimated by decision trees are also provided. Grey highlights taxa that have been domesticated

| Genus | Domesticated | Logistic regression | Tree all | Tree organismal |
| --- | --- | --- | --- | --- |
| Vicugna | Yes | 0.35 | 0.25 | 0.12 |
| Antilocapra | No | 0.35 | 0.09 | 0.33 |
| Ovibos | No | 0.35 | 0.25 | 0.33 |
| Capra | Yes | 0.35 | 0.89 | 0.33 |
| Ovis | Yes | 0.35 | 0.89 | 0.33 |
| Saiga | No | 0.31 | 0.09 | 0.33 |
| Equus (donkey) | Yes | 0.3 | 0.25 | 0.8 |
| Lama | Yes | 0.3 | 0.89 | 0.33 |
| Camelus (Bactrian) | Yes | 0.3 | 0.09 | 0.33 |
| Hemitragus | No | 0.3 | 0.89 | 0.33 |
| Pseudois | No | 0.3 | 0.09 | 0.33 |
| Ammotragus | No | 0.3 | 0.25 | 0.33 |
| Redunca | No | 0.29 | 0.05 | 0.2 |
| Budorcas | No | 0.28 | 0.09 | 0.33 |
| Equus (horse) | Yes | 0.27 | 0.89 | 0.8 |
| Equus (zebra) | No | 0.27 | 0.05 | 0.12 |
| Bison | No | 0.27 | 0.09 | 0.12 |
| Capricornis | No | 0.27 | 0.05 | 0.33 |
| Connochaetes | No | 0.26 | 0.05 | 0.12 |
| Cervus | No | 0.25 | 0.09 | 0.04 |
| Syncerus | No | 0.25 | 0.05 | 0.8 |
| Bos (taurine) | Yes | 0.25 | 0.89 | 0.8 |
| Bos (yak) | Yes | 0.25 | 0.89 | 0.14 |
| Kobus | No | 0.25 | 0.05 | 0.2 |
| Aepyceros | No | 0.25 | 0.05 | 0.33 |
| Oreamnos | No | 0.25 | 0.09 | 0.14 |
| Bubalus | Yes | 0.24 | 0.05 | 0.2 |
| Bos (zebu) | Yes | 0.24 | 0.05 | 0.8 |
| Hydrochoerus | No | 0.24 | 0.05 | 0.2 |
| Phacochoerus | No | 0.22 | 0.05 | 0.12 |
| Camelus (Dromedary) | Yes | 0.22 | 0.89 | 0.33 |
| Damaliscus | No | 0.22 | 0.05 | 0.12 |
| Hippotragus | No | 0.21 | 0.05 | 0.12 |
| Taurotragus | No | 0.2 | 0.05 | 0.14 |
| Oryx | No | 0.18 | 0.05 | 0.14 |
| Alcelaphus | No | 0.18 | 0.05 | 0.12 |
| Hylochoerus | No | 0.17 | 0.05 | 0.04 |
| Axis | No | 0.17 | 0.05 | 0.04 |
| Boselaphus | No | 0.17 | 0.05 | 0.14 |
| Myotragus | No | 0.15 | 0.25 | 0.33 |
| Potamochoerus | No | 0.14 | 0.05 | 0.04 |
| Hippocamelus | No | 0.14 | 0.09 | 0.04 |
| Pseudoryx | No | 0.14 | 0.05 | 0.2 |
| Litocranius | No | 0.13 | 0.05 | 0.04 |
| Pan | No | 0.12 | 0.05 | 0.04 |
| Sus | Yes | 0.12 | 0.89 | 0.04 |
| Tragelaphus | No | 0.12 | 0.05 | 0.04 |
| Beatragus | No | 0.12 | 0.05 | 0.14 |
| Rangifer | No | 0.11 | 0.09 | 0.04 |
| Cephalophus | No | 0.11 | 0.05 | 0.04 |
| Pongo | No | 0.1 | 0.05 | 0.04 |
| Addax | No | 0.1 | 0.05 | 0.14 |
| Macropus | No | 0.09 | 0.05 | 0.04 |
| Tapirus | No | 0.09 | 0.05 | 0.04 |
| Dicerorhinus | No | 0.09 | 0.05 | 0.04 |
| Dama | No | 0.09 | 0.25 | 0.04 |
| Blastocerus | No | 0.09 | 0.05 | 0.04 |
| Alces | No | 0.09 | 0.25 | 0.04 |
| Hippo (Madagascar) | No | 0.09 | 0.05 | 0.04 |
| Hexaprotodon | No | 0.09 | 0.05 | 0.04 |
| Gorilla | No | 0.08 | 0.05 | 0.04 |
| Rusa | No | 0.08 | 0.05 | 0.04 |
| Okapia | No | 0.07 | 0.05 | 0.04 |
| Odocoileus | No | 0.07 | 0.09 | 0.04 |
| Babyrousa | No | 0.06 | 0.05 | 0.04 |
| Elaphurus | No | 0.06 | 0.05 | 0.04 |
| Hippo (Crete) | No | 0.06 | 0.25 | 0.04 |
| Rucervus | No | 0.05 | 0.05 | 0.04 |

We can see that out of non-domesticated taxa *Antilocapra* and *Ovibos* come the highest on the list. Both are North American taxa, which hints that non-domestication of those animals might have had more to do with the place than the animals themselves. Quite close comes Saiga, which is a mysterious animal of semideserts, steppes and grasslands. Today they are critically endangered and found only in central Asian steppes. Saigas have been hunted for a long time, no idea why they have not been domesticated.

The next non-domesticated cohort includes *Hemitragus*, *Pseudois*, *Ammotragus, Budorcas* and *Redunca*. All except for *Redunca* are close taxonomic relatives of goats and sheep, and all are from Palearctic. Since goats and sheep have already been domesticated at the same region, perhaps there was no strong need to go for a similar wild option rather than choosing to breed already domesticated species.

Of those domesticated *Bos* and *Bubalus* come rather low in the ranking, maybe because they are relatively more specialized and sensitive to food quality than the higher cohort including sheep, goat, camelids and horses.

*Sus* has been domesticated by comes very low in the ranking. *Sus* is an omnivore having their diet relatively close to that of humans, and it is indeed somewhat surprising that multiple domestications of *Sus* happened.

### 3.3. Realized and potential rates of domestication across biogeographic realms

Now that we have estimates for suitability for domestication, next we analyse how potential suitability is distributed across biogeographic realms, and how does it relate to observed rates of domestication across the realms. Table 5 gives the rates for 10 domesticated genera. In addition, Figure 3 plots potential domestication rates across all the ranking of candidate genera, first assuming that the genus with the highest estimate is domesticated, then that two genera with the highest scores are domesticated and onwards.

**Table 5.**
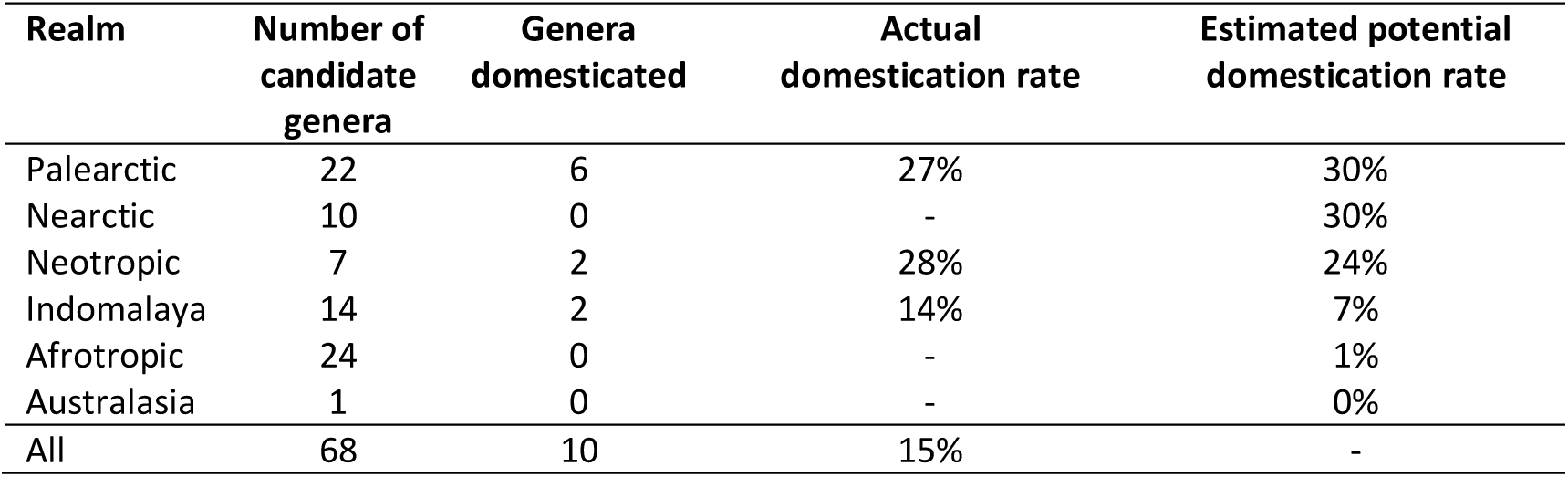
Actual and estimated potential rates of domestication across biogeographic realms. In some realms several species or subspecies of the same genera were domesticated (e.g. *Bos*), in such a case a genus is counted only once. Given potential rate of domestication is *assuming* that top-10 genera with highest estimated propensity for domestication are domesticated. The last row (All) is not a sum of all other rows, because some genera occur in several realms.

| Realm | Number of candidate genera | Genera domesticated | Actual domestication rate | Estimated potential domestication rate |
| --- | --- | --- | --- | --- |
| Paleartic | 22 | 6 | 27% | 30% |
| Nearctic | 10 | 0 | - | 30% |
| Neotropic | 7 | 2 | 28% | 24% |
| Indomalaya | 14 | 2 | 14% | 7% |
| Afrotropic | 24 | 0 | - | 1% |
| Australasia | 1 | 0 | - | 0% |
| All | 68 | 10 | 15% | - |

**Figure 3.**
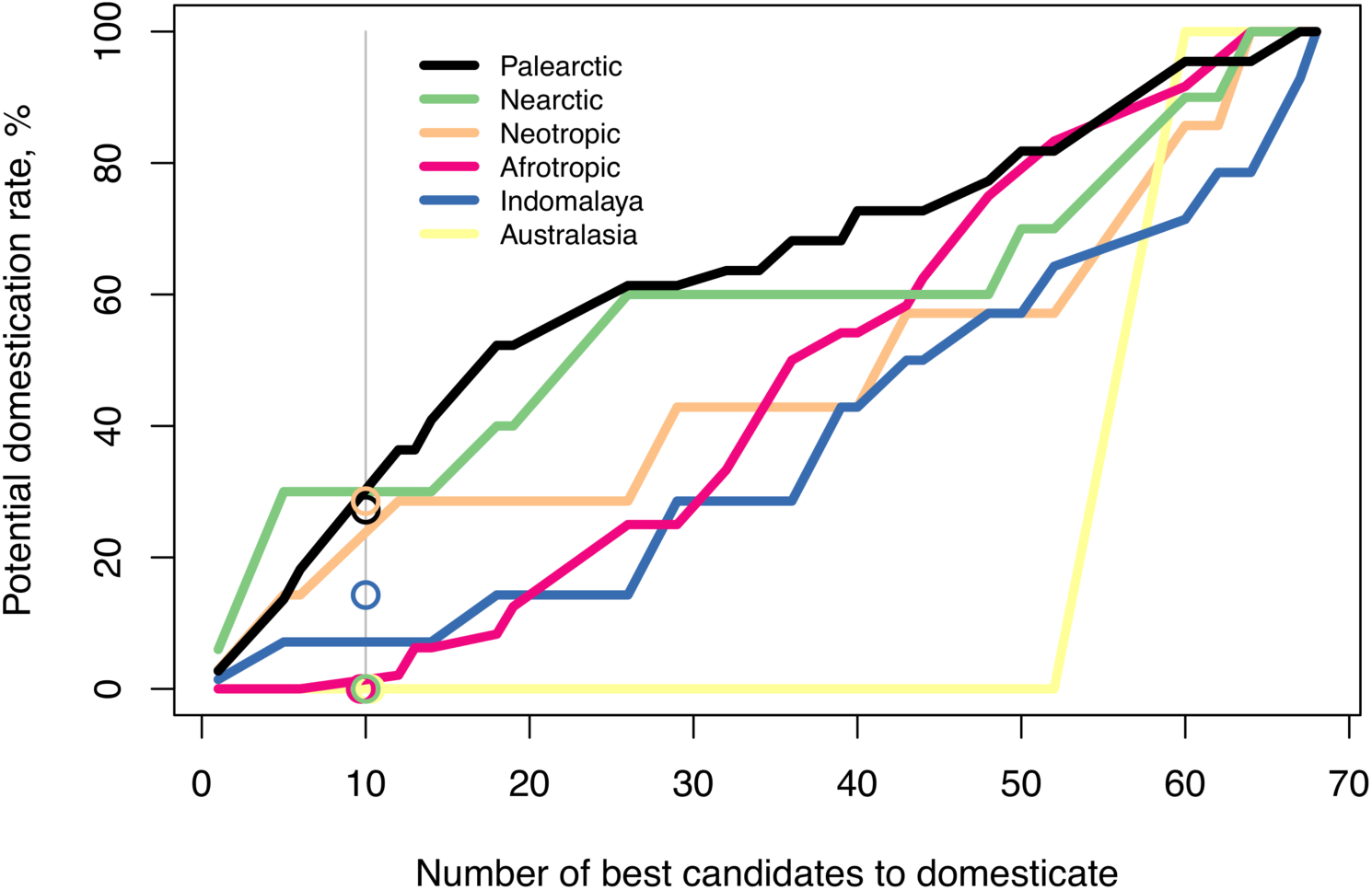
Potential domestication rates considering different number of best candidates. Circles denote real observed rates of domestication. The vertical line denotes the number of genera that were actually domesticated. The horizontal lines are derived from estimated ranking of genera for domestication.

From the vertical alignment of lines in Figure 3 we can see three groups emerging: Palearctic, Nearctic and Neotropic seem to host the largest share of suitable genera (the highest vertical position), followed by Afrotropic together with Indomalaya, and finally, Australasia hosts the least. This can be interpreted as no matter how many candidates each realm hosts, how suitable their candidates are on average. Interestingly, while the logistic regression model given in Equation (1) used for estimating suitability scores does not include latitudinal information (“tropical”), latitudinal patterns do emerge. Temperate realms seem to host proportionally roughly 4 times more suitable taxa than tropical. Neotropic realm includes is a mix of tropical and non-tropical environments, and appears in-between of the temperate and tropical groups. Australia has only one genus large enough as a candidate and it gets a low estimate, thus Australia lags quite behind from both groups.

Our analysis suggests that, Australia discarded, the potential suitability was surprisingly consistent across the biogeographic realms: around 30% in temperate, less than 10% in tropical, and around 20% in the Central and South America, which is a mix of temperate and tropical.

Most interesting is the relation between actual and potential domestication rates. In Figure 3 we see that Palearctic and Neotropic have their actual rates (circles) quite near to their potential rates, and so does Australasia and Africa. Two exceptions appear: Indomalaya seems to have higher actual rates than the potential, and Nearctic (North America) has zero actual rate, while the potential appears to be as high as that of Palearctic.

Diamond (1997) argued that uneven rates of domestication across the globe were primarily due to animal characteristics rather than geographic location, which from our analysis seems to be only partially true. It is noteworthy that Diamond (1997) counted domestication at the species, and analysed continents putting together South and North America, and not distinguishing Indomalaya, which is climatically very different from the rest of Asia. He got 18% domestication rate in Eurasia, 4% in the Americas and 0 elsewhere. Our analysis taking into account biogeographic realms suggests that while the total number of candidates in South America was very small, South America (Neotropic) had very similar actual domestication rates to Eurasia (Palearctic). North America (Nearctic), on the other hand, had seemingly many candidates, but actual domestications did not happen.

The distribution of actual and potential rates of domestication across the realms suggest that while each realm consistently hosted a pool of suitable animals, other factors than just suitability of animals themselves must have been driving domestication rates, but at the moment we have no explanation for the large North America mismatch between actual and potential. As for availability for availability of potentially suitable livestock animals, we hypothesize that a large part of explanation might be biogeographic, requiring a particular intersection of cold and hot semi-arid climatic zones. In the following section we discuss evidence for this along with our interpretations.

## 4. BIOGEOGRAPHIC ANALYSIS

To explain suitability scores for domestication across biogeographic realms, we suggest an interplay between climatic zones primarily suitable for plant domestication and animal domestication. The argument goes as follows: hot Semi-arid climate was most suitable for plant domestication, cold Semi-arid climate selected for animals most suitable for domestication as livestock, the realized rates of domestication, with a major exception of North America, reflect the how much intersection of hot and cold Semi-arid climatic zones was available at each continent.

### 4.1 Biogeography of plant domestication

Sedentary lifestyle started with plant domestications at least half a millennium earlier than the first known livestock animal domestications (Larson et al 2014). Many arguments have been put forward why domestication of plants started at this particular time. Early arguments, such as the Oasis hypothesis (Childe 1928), favoured climate change. Recently more emphasis has been put on socioanthropological causes, such as depletion of wild food resources (Diamond 1997). Undoubtedly reasons for the time and place were multiple, among which was the warm seasonal climate at the Fertile Crescent. This climate favoured a bimodal way of life for plants and animals: feasting during the rich seasons, and being dormant during the lean seasons. Such a bimodal operation requires effective storage of nutrients, as well as fast onset and growth at the start of the rain season. Plants can cope with seasonal aridity, among other ways, by producing large protein-rich seeds (Diamond 1997), that are nutritious and preserve well. Such seeds make a nutritious storable material, considered to be favourable for domestication.

Assuming that climate stress selects for domestication-friendly plant material, the next question is where in the world such climates could be found at the times of early domestication. Based on climate of the Fertile Crescent today, Diamond (1997) argued that Mediterranean climate presents the most favourable climatic conditions for plant domestication. In the Koppen climate classification system (Kottek 2006) Mediterranean climate is characterized by dry summers and mild wet winters with the average temperature of the coolest month 0-22°C and the cumulative precipitation in the driest summer month less than 30-40 mm. Figure 4a depicts the areas of Mediterranean climate today. Diamond (1997) argued that out of all the Mediterranean zones the Fertile Crescent was favourable because it was the most inland.

**Figure 4.**
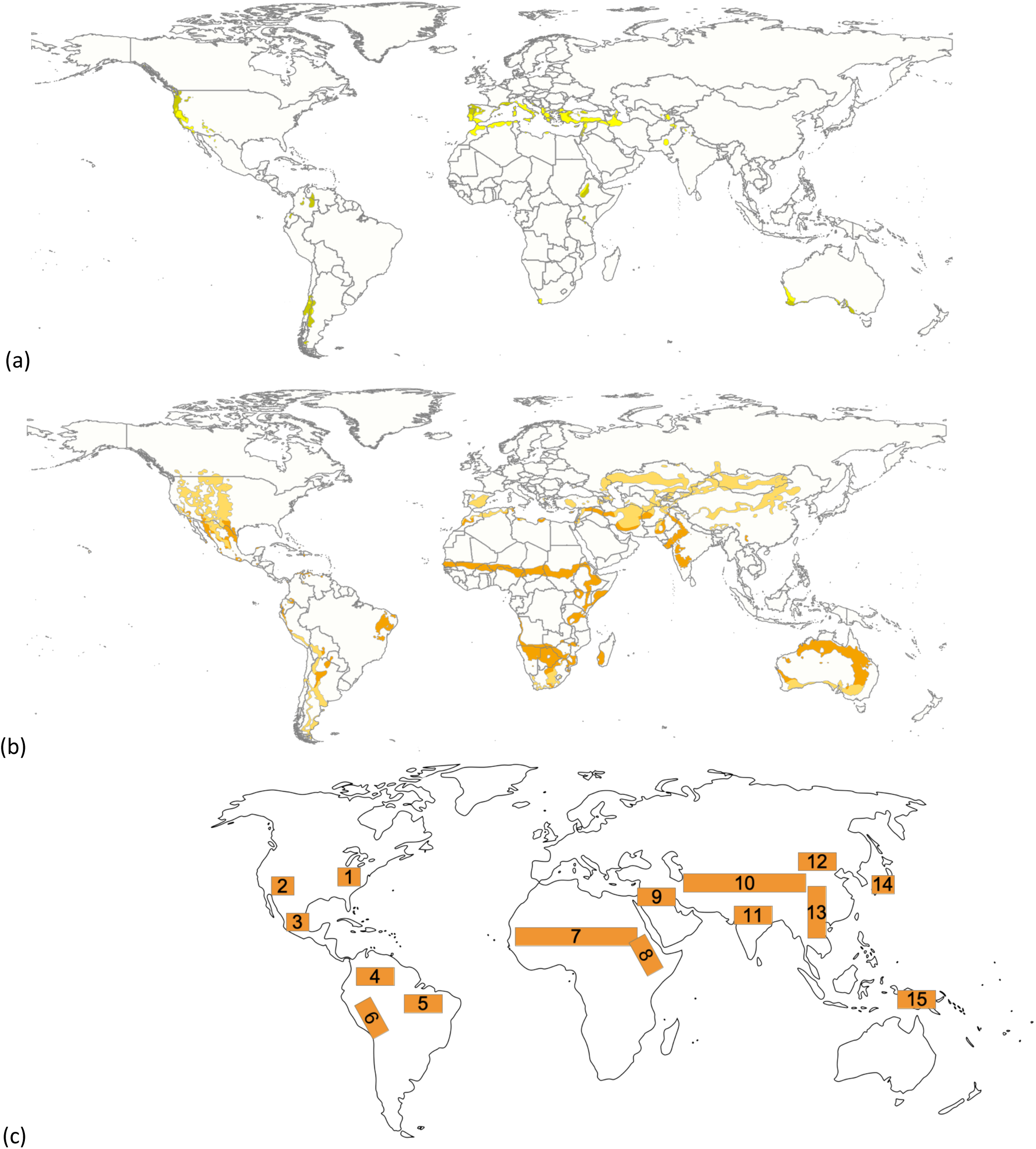
Mediterranean and semi-arid climate zones today by Koppen climate classes. (a) Mediterranean climate zones today (light yellow – Csa, hot-summer Mediterranean climate; dark yellow - Csb, warm-summer Mediterranean climate). (b) Semi-arid (steppe) climate zones today (dark yellow – BSh, hot semi-arid zone; light yellow – BSk, cold semi-arid zone). In dark yellow it does not freeze, in light yellow the coldest month has freezing temperatures. (c)Areas of domestication of plants and animals with examples for each centre: (1) Eastern North America: chenopodia, squash, sunflower, and maygrass; (2) Southwest US: turkeys; (3) Mesoamerica: maize, squash, beans, and turkeys; (4) northern Peru/Ecuador: squash and lima beans; (5) Amazonia: manioc, yams, peanuts, and Muscovy duck; (6) Andes: oca, potato, quinoa, amaranth, llama, alpaca, and guinea pigs; (7) sub-Saharan Africa: pearl millet, sorghum, and African rice; (8) Horn of Africa/Nile Valley: asses, tef; (9) Near East: wheat, barley, lentils, peas, sheep, goats, taurine cattle, and pigs; (10) Central Asia: horses, golden hamster; (11) South Asia: browntop millet, water buffalo, and zebu cattle; (12) North China: foxtail and broomcorn millet; (13) South China/Southeast Asia: rice and chickens; (14) Japan: barnyard millet, mung bean, burdock; (15) New Guinea: bananas, yams, and taro. The list of domesticates not exhaustive. Map (c) is drawn after Zeder (2017).

Mediterranean climate zones today are ideal for wine production. This climate primarily generates scrubby and dense vegetation – broad-leaved evergreen shrubs, bushes and small trees. Plants of this kind are not ideal targets for plant domestication, not only because it takes several years for first fruits to appear. Such a vegetation cover would have been tiresome grounds for early agriculture, because the land would continuously have needed to be cleared of bushes and shrubs. Slightly harsher (dryer) climate, on the other hand, would have kept large parts of land clear from woody vegetation naturally.

Indeed, the first domesticated plants were grasses (wheat and barley). It is conceivable that the climate could have been harsher than the Mediterranean to select for grasses over woody shrubs. The next zone towards harshness is the Semi-arid zone, characterized by low and very seasonal annual precipitation. The precipitation threshold for Semi-arid zones depends on whether the main precipitation comes in winter or in summer and how it interplays with the temperature and the length of daylight. Vegetation in these zones is short or scrubby, dominated by grasses or small shrubs. The coverage is depicted in Figure 4b.

Visually, Semi-arid zones in Figure 4b coincide with many areas of early agriculture, drawn after Zeder (2017) in Figure 4c: The Fertile Crescent (#9), sub-Sahara (#7), Nile Valley (#8), India (#11), North China (#12), Mesoamerica (#3), Andes (#6). All these areas domesticated grass plants or herbaceous plants.

Several areas of early plant domestication are outside the proposed Semi-arid zone pattern, in particular – rise domesticated in South China, bananas, yams and taro in New Guinea, millet, bean and burdock in Japan, as well as some parts of Americas. Of those, rice from China and bananas from New Guinea are perhaps most widely cultivated today. Both places have quite seasonal (New Guinea in highlands), but not particularly arid climate.

On the other hand, two hot Semi-arid areas, Kalahari in Africa and Australian coastlands, hosted no known plant domestications. Despite these open questions global match between Semi-arid climatic zones and areas of early plant domestication is quite striking, suggesting that hot Semi-arid zones denoted as BSh in the Koppen climate classification system (Kottek et al 2006) rather than Mediterranean zones (CSa), might have the primary climate for onset of agriculture.

### 4.2 Cold seasonal environments select for suitable livestock animals

Transitioning to a sedentary lifestyle and plant domestication required early humans to settle in places suitable for agriculture. In such circumstances humans who have settled for agriculture would naturally look for domesticable animals nearby. Diamond (1997) pointed out a curious pattern: while the first livestock animals were domesticated at the Fertile Crescent, all of them presumably evolved in Central Asia, suggesting initial adaptation to much colder seasonal environments. And indeed, our computational analysis reported in the previous section hinted that seasonal semi-arid zones, such as cold steppe, are likely to host animals most suitable for livestock domestication.

Strong seasonality prescribes two modes of existence: the prosperous season during which the ecosystem is flourishing, and the lean season during which the ecosystem shuts down to bare essentials. While small mammals can sleep or hide (Liow et al 2009) through the bad season, large mammals living in seasonal environments have little alternatives but to migrate in order to survive the lean season. For successful migration, it is critical for the group to follow the leader (Guttal and Couzin 2010), as commonly observed, for example, in birds. Therefore, seasonal environments must select for follow-the-leader social structure, particularly in large mammals. Alternation between seasons requires accumulation of energy during the rich season in order to survive the lean season. Therefore, seasonal environments must also select for rapid growth of offsprings to be ready to realocate when time comes, and overall rapid growth of body mass in order to accumulate resources for the lean season.

Hypsodonty, which characterises durability of teeth and tolerance of abrasive food, appears to be one of the strongest explanatory variables for domestication in our analysis. Food for domestic animals needs to be common in human habitats and available in bulk, since specialized diets such as meat, fish, nuts, fruits or aquatic plants would take a lot of effort to collect and would not be economically viable. More importantly, the diet of domestic animals needs to be different from preferred human foods, since otherwise a more energetically efficient solution for humans would be to eat that food themselves. Grass is available in bulk and has low requirements for maintenance in semi-arid climatic zones and humans cannot utilize it directly due to lack of ability to digest cellulose. Not surprisingly, most of livestock animals appear to be mixed-feeders that given hypsodonty and digestive specializations can tolerate or even prefer grass.

Fast growth might be associated with seasonality and migration. Animals experience seasonality in both tropical and temperate environments, but in our analysis tropical habitats have a strong negative association with domestication. Indeed, a curious question is why some tropical groups, such as antelopes, have never been successfully domesticated. Since the tropical savanna animals have been hunted for a long time, they must be at least eatable for meat. Many African antelopes come from marginal and very seasonal environments and presumably have similar migratory and growth strategies for coping with the bad season, but hot rather than cold.

The main difference between the two environments is that in cold seasonal environments temperature is the limiting factor, while in hot seasonal environments seasonal lack of precipitation is the limit. Thus, cold seasonal environments are likely to select for fast growth of fat to help to withstand changing temperatures, while tropical seasonal savannas are likely to prioritize selection for minimum loss of water. Indeed, most of grazing antelopes native to such conditions, such as Oryx, can survive without drinking water for long periods^3^. Indeed, in our analysis water dependency is among the main explanatory variables for probability of domestication along with hypsodonty, migration, and non-tropical habitats.

For domestication to be successful animals should have an incentive to stay domesticated. An animal could escape predators for years, survive without food for weeks, but a water-dependent animal would hardly survive without water for more than a couple of days. Since humans are water-dependent themselves, they are always in reach of drinking water, and may have technology for finding and storing water. That could be an incentive for animals that regularly need drinking water to stay close to humans.

Some other variables in our analysis, such as migration, grazing, and highlands have the strongest association with domestication in the middle of their value range. Such animals can be characterized as “migratory, but not too much”, “grazer, but not to an extreme”, “highland habitats, but not too high up”. This links back to observations from analyses of evolutionary processes, suggesting that the frontier for speciation, so called “species factory” (Bernor et al 1996, Fortelius et al 2014), is often not at the extreme ends of the environment, but right behind the extreme frontier. Semi-arid environments which have produced most of material for domestication (plants and animals) are somewhat like this – they are close to deserts, but not yet deserts.

Following our computational analysis we propose that cold seasonal habitats, particularly alpine environments, are the highly suitable for selecting valuable candidates for potential domestication as livestock. Our analysis also showed that animals from all over the trait space have been domesticated, but those of marginal environments (such as goats, llamas, camel, sheep) were the most densely representing domestication in our character space. Animals from marginal environments do not have luxury to be highly specialized, they can efficiently process low quality vegetation of various kinds appear to be the most valuable, since they help to convert low quality fibrous vegetation, which humans otherwise would not utilize, into energy that humans can use.

### 4.3 Crossroads at the Fertile Crescent

While hot Semi-arid climate zones in Figure 4b appear to be the most common for plant domestication, cold Semi-arid zones highlighted in the same figure appear to be common to produce animals highly suited for domestication. We propose that places where hot semi-arid and cold semi-arid zones meet have been the most suitable for early sedentary life. Indeed, the Fertile Crescent is one of several such zones. While the match in Figure 4b is based on Semi-arid climatic zones today, the final question for our analysis is how such zones were distributed around 10 000 years ago at the time of first domestications, and whether their global distribution can potentially explain different rates of domestication across continents.

The best currently accessible global climatic map of the past depicts vegetation at the Last Glacial Maximum, that is 15 000 – 25 000 years before present (Ray and Adams 2001). Figure 5a depicts our projection of Ray and Adams (2001) climate reconstruction to the Koppen climatic classes, and Figure 5b depicts Semi-arid zones today (Kottek et al 2006) along with deserts for a direct comparison. At the time of early domestication of plants and animals, about 10 000 years before present, geographic distribution of the climatic zones must have looked as an interpolation of Figures 5a and 5b. Judging from these maps, the most notable changes in the hot Semi-arid areas have occurred in the Fertile Crescent, North-East Africa and Central America. In those locations hot Semi-arid areas were more widely spread than today. Two other major changes of Semi-arid areas have occurred in South America and Australia. In South America cold Semi-arid areas have shrunk, and in Australia hot Semi-Arid areas have expanded.

**Figure 5.**
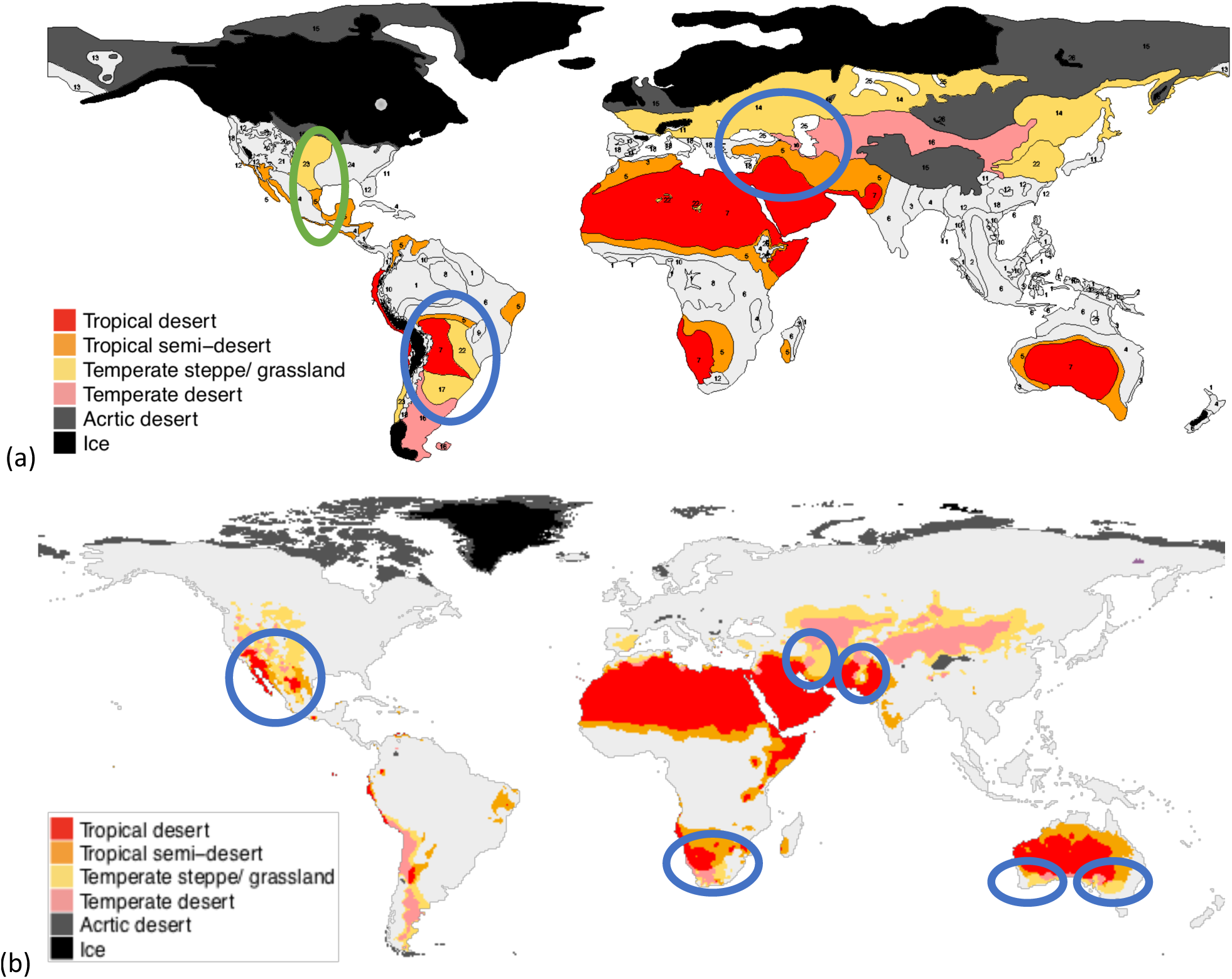
Climate zones in the past and today. (a) Vegetation cover at the Last Glacial Maximum (15 000-25 000), the plot is adapted from Ray and Adams (2001). (b) Desert and Semi-arid climate zones today (Koppen climate class B). Blue ellipses indicate intersections of hot and cold Semi-arid zones with arid zones nearby. Green ellipse indicates the are without apparent more extreme zones nearby.

Looking at the distribution of zones around 20 000 years ago (Figure 5b), only two zones, marked by blue ellipses, appear to provide large enough interfaces between hot and cold marginal environments. By and large it seems that the Fertile Crescent and Central South America areas were the largest interfaces of suitable environments, and those are the continents where the highest rates of domestication occurred.

Climatically speaking, temperate desert is mainly formed in Central Europe and North America due to topographic effect and monsoon climate^4^. Monsoon climate makes the summer drier in Central Eurasia, Arabian Peninsula and Mediterranean. These regions are mostly covered by cold steppe climatic zone (Bsk). North America does not have much land in sub-tropical region, so there is little room for subtropical desert there. Only Eurasia and Africa have large areas of subtropical arid area. As a result, Arabia becomes the major region in which the two climatic zones meet.

North America also seems to have had a reasonably large suitable interface, but at least visually the extreme frontier (hot or cold deserts) are lacking in Figure 5a, making the North American Semi-arid areas (highlighted in green). Yet, at the present day extreme areas are present, thus, lack of domestication in North America remains an open question, particularly given that our analysis has identified about the same average suitability of species as in Eurasia (Palearctic). Indomalaya domestications are few, and do not quite fit the common pattern of cold environments. Perhaps humid and hot tropical settings required locally adapted animals, and the two domesticated ones -- *Bubalus* and *Bos* (zebu), were the most suitable locally.

## 5. CONCLUDING REMARKS

Human society today relies on a small number of livestock animals, and we keep wondering what made those taxa particularly suitable, was it about the animals themselves, the environments, the geography, the timing, or perhaps an interplay of all. As Alfred Crosby (2006) phrased it, domestication was often discontinuous and did not always work. The patterns we have analysed are results of what has worked.

Our analysis showed that the most suitable animals for domestication were generalists adapted to persistence in marginal environments of low productivity, largely corresponding to cold semi-arid climate zones. Such animals would have inhabited temperate steppe or grasslands. At the time of first domestications the global distribution of such areas should have been somewhat interim version in between of map 5a and map 5b. Yet by far the largest, and almost the only area for such animals to come from is likely to have been Central Asia, and that is where the big four domestic animals are thought to have originated from. The Fertile Crescent has been and is the major intersection of hot and cold semi-arid climate zones. If hot semi-arid zones select for suitable plants, cold semi-arid zones select for suitable animals, an intersection of the two zones must be an ideal place to domesticate both kinds.

Many questions are still open, one of which is why animals from marginal hot environments apparently are unsuitable for domestication. Even if animals from cold environments might be preferable due to growth rates, growth rates do not vary that much given the body mass, and this should automatically disqualify animals from hot climates. The Sahara desert is not very likely to have been a barrier either. The difference must be related to adaptations to the tropical habitats. We have hypothesized that water dependency and predator density are important for the follow-the-leader social structure, but some of the hot arid climate grazing antelopes, like waterbuck, are water dependent, and some, like African buffalo, are quite resistant to predators. We hope for interesting studies to follow.

## Acknowledgements

I thank Mikael Fortelius, Jared Diamond, Marcus Clauss, Miikka Tallavaara and Hui Tang for discussing early versions of this study. Any errors, omissions or misinterpretations are my own. An earlier version of this study was submitted to and rejected from Environmental Archaeology journal. I thank three anonymous reviewers for their feedback. I did incorporate many of the suggestions and I did my best to clarify the points that were misunderstood. I may or may not to resubmit to another journal, for now I am happy to make the study available via bioRxiv. This is a contribution from the Valio Armas Korvenkontio Unit of Dental Anatomy in Relation to Evolutionary Theory.

## SUPPLEMENT

### A.1 Domestication of the first known livestock animals

To the best knowledge today, goat and sheep were the first herbivorous animals to be domesticated about 11000 years ago in different parts of the Fertile Crescent (Zeder 2008). Domestic sheep today are grazers, while domestic goats are browse-dominated mixed-feeders. The nearest wild relatives of both – mouflon sheep (*Ovis orientalis*) and wild goat (*Capra aegagrus*) have numerous subspecies, most of which live in mountain habitats and are grazers or graze dominated mixed-feeders. Mouflon sheep are very migratory, in winter they reallocate to lower altitudes. Most of wild goat subspecies are extremely endangered, found in small areas. At least Ibex is reported to be migratory, relocate to alpine pastures in spring. Due to their habitats both species are difficult to reach for predators (except human hunters). Both are reported to be grass-dominated mixed feeders. Both need drinking water daily in summer season.

Pigs were first domesticated about 10500 years ago in the Fertile Crescent, and soon afterwards independently in China. Pigs are omnivorous, which allows broad diet. Apparat from pigs the only other large omnivorous mammals are bears, but they are not migratory, they hibernate. The nearest wild relative of domestic pig is wild boar (*Sus scrofa*), native to much of Eurasia and North Africa. There are many subspecies. Boars are social animals, living in female-dominated groups. While the main habitats are deciduous forests, wild boar occupies a diverse range of habitats from taigas to deserts, as well as alpine zones, as long as there is no regular snowfall. They are highly water dependent and they are migratory, especially along elevation gradient.

Cattle were first domesticated around 10000 years ago in the Fertile Crescent from wild aurochs (*Bos primigenius*), which are now extinct. Eurasian aurochs ranged in Europe, Siberia and Central Asia, and East Asia. Aurochs formed herds (van Vuure 2005). Auroch is thought to have been a grazer or mixed-feeder. The original habitats are uncertain, but it is thought to have lived last years of existence in wet forests or marshlands. Isotopic analysis of aurochs from Denmark suggests a habitat/dietary shift of wild aurochs from open grasslands to forests and marshes after domestic cattle arrived to Denmark (Nygaard et al 2005). Aurochs had fur.

Table A1 lists assumed ancestor species of the present livestock animals.

**Table A1.** Assumed wild ancestors of the livestock animals.

| Domestic animals |
| --- |
| Sheep (*Ovis aries*) |
| Goat (*Capra aegagrus hircus*) |
| Pig (*Sus scrofa domesticus*) |
| Cattle (*Bos taurus / indicus*) |
| Horse (*Equus ferus caballus*) |
| Donkey (*Equus asinus*) |
| Llama (*Lama glama*) |
| Alpaca (*Vicugna pacos*) |
| Water buffalo (*Bubalus bubalis*) |
| Yak (*Bos grunniens*) |
| Camel (*Camelus bactrianus*) |
| Camel (*Camelus dromedaries*) |

#### A.2 Scoring scheme for habitat, dietary characteristics and dental traits

All the variables except for hypsodonty are encoded as categorical in the scale from 0 to 1, such that the variables could potentially be used as numeric in the analysis. A detailed description of the scoring scheme is given in the Supplement.

##### 1. Grazer

Encoding: 1 means grazer, 0.67 encodes graze-dominated mixed-feeder, 0.33 encodes mixed-feeder that takes a small portion of grass, for instance, browse-dominated mixed-feeders are encoded this way, 0 means non-grazer, and mostly includes browsers, also frugivores and omnivores may appear in this category.

##### 2. Highly specialized

This variable is meant to distinguish regular grazing-browsing herbivores from specialized diets, such as aquatic plants or fruits. Encoding: 0 encodes regular grazing or browsing herbivore, 1 encodes highly specialized diets.

##### 3. Water dependence

This variable primarily means dependence on drinking water. Encoding: 1 means the animal needs to drink daily, 0.5 means drinking every 2-3 days, 0 means the animal can go a week or more without drinking water. The last category includes aridity adapted animals that do not need to drink water at all (e.g. *Oryx*), they get their water with food. Category 1 also includes aquatic animals, such as *Hippopotamus* or *Bubalus*.

##### 4. Open habitat

Encoding: 1 grassland, steppe, open savanna, semi-desert, mountain terrain; 0.5 woody savanna, bushlands, sparse forest; 1 dense forest.

##### 5. Highland habitat

This is meant to encode habitats at high elevation, mountainous or hilly terrains. Encoding: 1 mountains or hills, 0.5 foothills, 0 steppes or forests, more or less flat terrain.

##### 6. Migration

This variable is meant to encode seasonal migration. Historical migration information was considered, where possible, for example, *Bubalus* or *Bison* have been migratory in the past and therefore are scored as migratory. Encoding: 1 regular seasonal migration to different geographic areas, 0.5 occasional seasonal migration or seasonal expansion/decline in range, 0 no migration.

##### 7. Highly endemic

This variable is meant to describe genera occurring in very restricted geographic areas, typically islands. Encoding: 1 highly endemic, 0 wide distribution.

##### 8. Tropical

This variable is meant to capture biogeographic distribution, following biogeographic realms (Figure A1). Encoding: all genera from Afrotropic and Indomalaya realms are assigned to category 1 – tropical, all genera from Palearctic and Nearctic are assigned to category 0 – non-tropical, genera from Neotropic and Australasia are assigned to one or the other on case-by-case basis.

Dental trait scoring generally follows Zliobaite et al (2016). We leave out horizodonty and cement, since they have shown relatively little global signal in the past analyses. Scoring is typically done on the second upper molar, except for suids, where functionally dominant tooth is the third upper molar. Dental trait scores come from publications (Zliobaite et al 2016, Galbrun et al 2018, Zliobaite et al 2018 and Oksanen et al 2018). *Pseudoryx* teeth are scored by the nearest living relative (Bubalus or Syncerus, as per Bibi 2013).

##### 9. Hypsodonty (HYP)

Hypsodonty refers to ordinated molar crown height: 3 is for hypsodonty, where the tooth height-to-length ration is roughly higher than 1.2, 2 is for mesodonty, where the height-to-length ratio is 0.8-1.2, and 1 is for brachydonty, where the ratio is less than 0.8. Figure A.1 gives examples of brachydont, mesodont and hypsodont traits.

**Figure A1.**
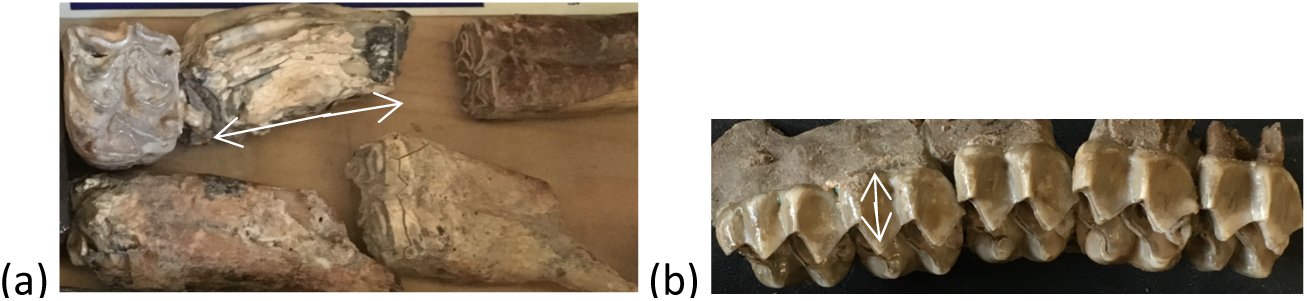
Examples of hypsodonty scores: (a) hypsodont (*Equus*), (b) brachydont (*Anchitherium*).

##### 10. Acute lophs (AL)

This variable indicates presence (or absence) of sharp-bladed cutting structures, typically extending across two or more cusps. A cusp is one developmental unit of a tooth, it typically has a pointy shape. Encoding: 1 acute lophs are present, 0 absent.

##### 11. Obtuse lophs (OL)

This variable indicates presence of blades, which are not sharp. Sharpness here is not determined by tooth wear (such as mesowear), but by how angled the blade is. Obtuse lophs generally are characterized by obtuse angles. Encoding: 1 obtuse lophs are present, 0 absent.

**Figure A2.**
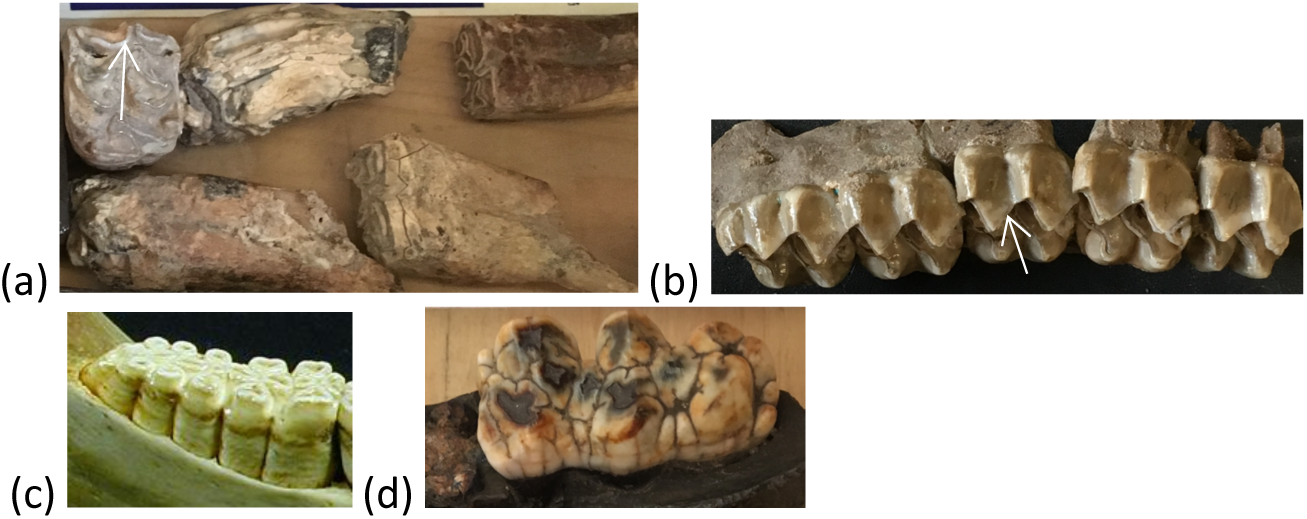
Examples of different AL and OL scores: (a) obtuse lophs present, acute lophs absent (*Equus*), (b) acute lops present, obtuse lophs absent (*Anchitherium*), (c) no lophs (*Phacochoerus*), (d) no lophs *(Sus*).

##### 12. Structural fortification (SF)

This variable encodes differential thickening of enamel that results in cusps remaining to stand out throughout tooth wear. Encoding: 1 structural fortification of cusps is present, 0 absent.

**Figure A3.**
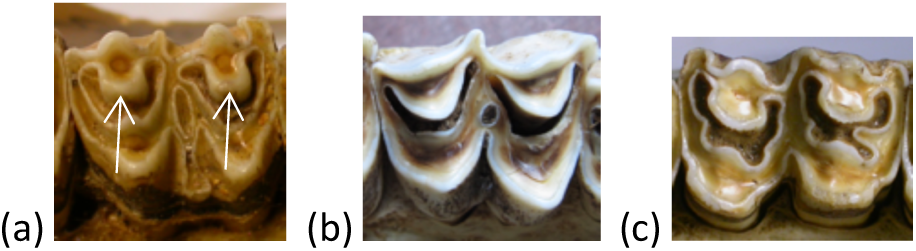
Illustration of SF: (a) Present (*Redunca*), (b) absent (*Tragelaphus*), (c) absent (*Alcelaphus*).

##### 13. Flatness/ occlusal topography (OT)

Encoding: 1 occlusal topography is flat, 0non-flat

**Figure A4.**
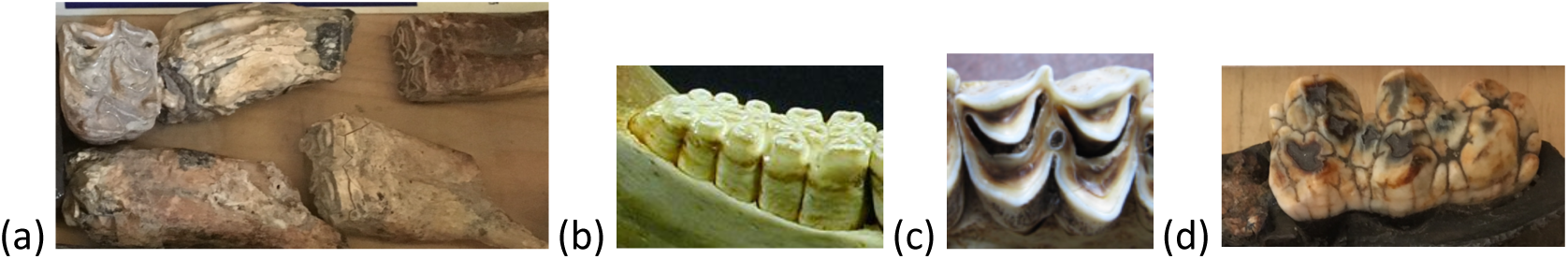
Examples of different OT scores: (a) flatness present (*Equus*), (b) present (*Phacochoerus*) (c) flatness absent/non-flat (*Tragelaphus*), (d) absent (*Sus*).

##### 14. Goat-like morphology (GO)

The purpose of this variable is to capture non-specialized herbivore dental morphology, where only obtuse lophs are present, but other specialized loph features, like acute lophs, structural fortification or flat occlusal topography are absent. This variable does not need to be scored, it can be computed from OL, AL, SF and OT. Encoding: 1 if OL =1 and AL=0 and SF = 0 and OT = 0, else 0.

**Figure A5.**
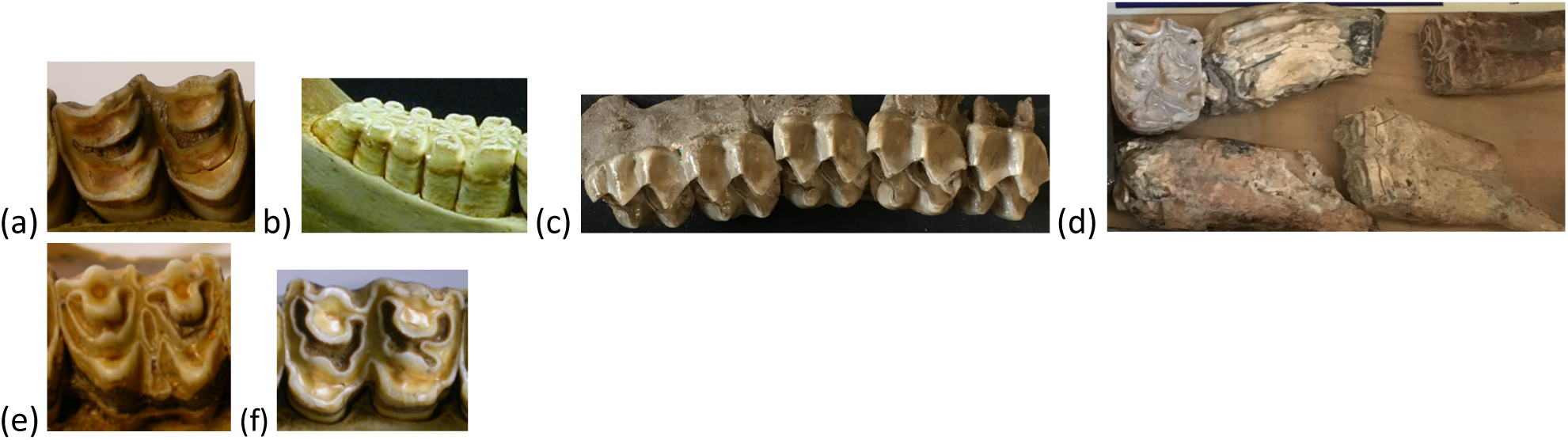
Examples of different GO scores: (a) present (*Antidorcas*), (b) absent due to no lophs and flat occlusal surface OT=1 (*Phacochoerus*), (c) absent due to acute lophs AL = 1 (*Anchitherium*), (d) absent due to flat occlusal surface OT = 1 (*Equus*), (e) absent due to structural fortification SF = 1 (*Redunca*), (f) absent due to flat occlusal surface OT = 1 (*Alcelaphus*).

##### 15. Bunodont (BU)

This variable encodes teeth that have no lophs, for example, human teeth. This variable does not need to be scored, it can be computed from OL, AL, SF, OT. Encoding: 1 if OL = 0 and AL = 0 and SF = 0 and OT = 0, otherwise 0.

**Figure A6.**
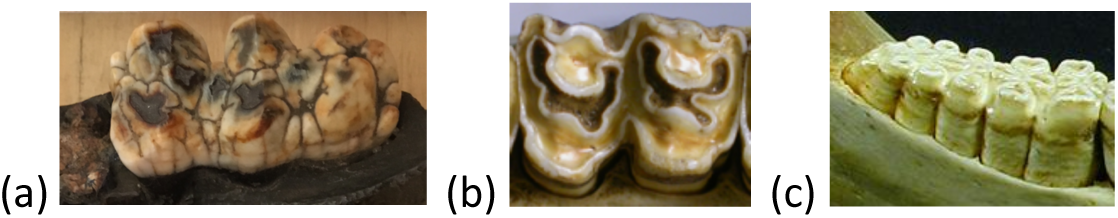
Examples of different BU scores: (a) bunodont (*Sus*), (b) non-bunodont, has lophs (*Alcelaphus*), (c) non-bunodont, due to flat occlusal surface (*Phacochoerus*).

The status of domestication is assigned following Diamond (1997), Larson et al (2014), and Zeder (2008). Even though domestication attempts on other taxa might have happen in the past, we label animals as domesticated based on the present day status, that indicates success in domestication of a given taxa.

#### A.4 Logistic regression fit on all 15 candidate variables

The resulting logistic regression model for the probability of domestication fit on all 15 candidate variables given in Table 2 is

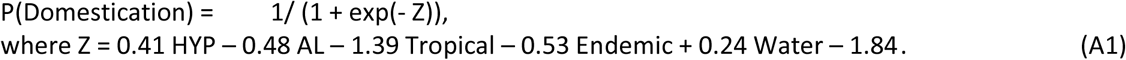

#### A.4 Other supplementary information

Figure A7 maps biogeographic realms, which we use as units of analysis for rates of domestication.

Figure A8 gives the linear correlations between the explanatory variables describing candidates for domestication.

Table A2 gives the dataset of candidate genera, and Table A3 lists genera that have been excluded from the candidate list.

Table A4 gives the first columns of the rotation matrix of the principle component analysis.

**Figure A7.**
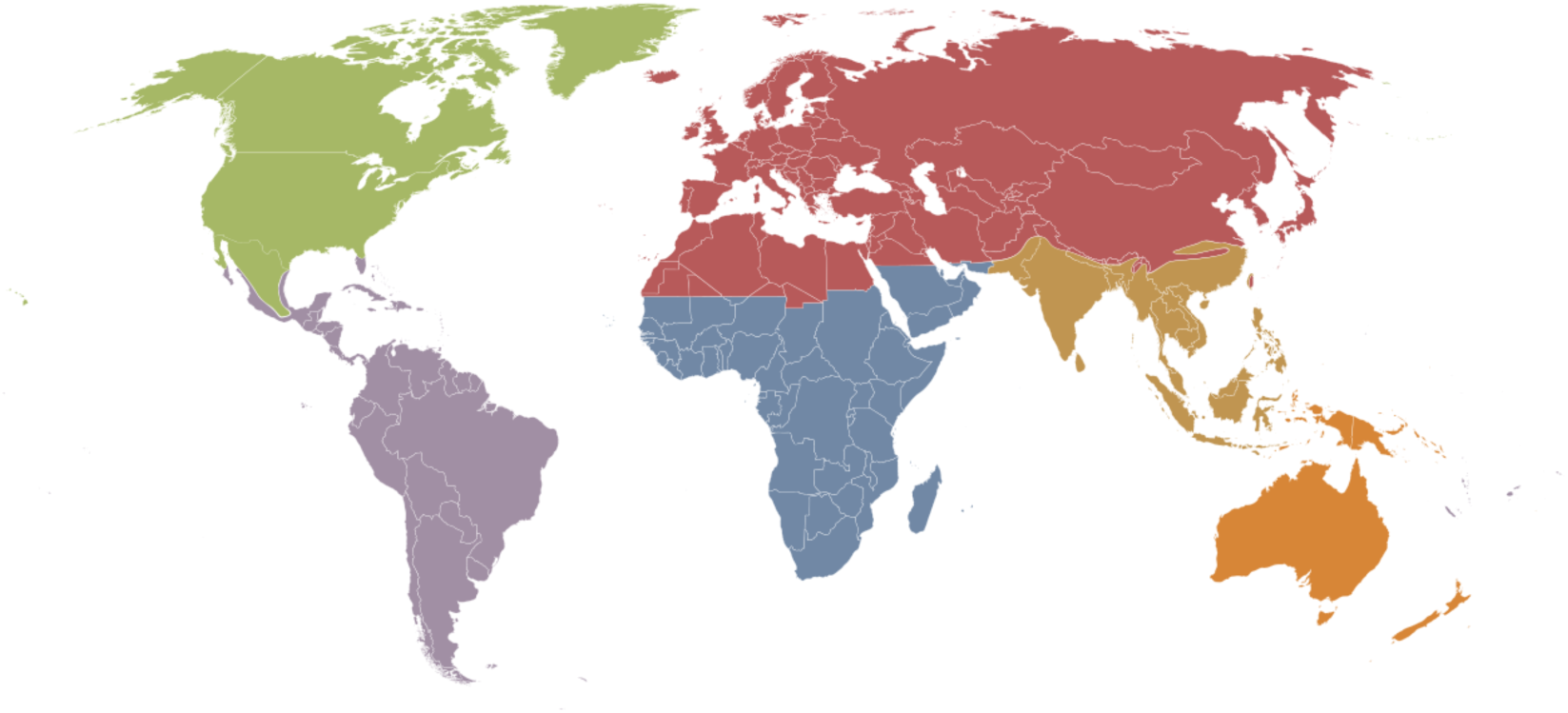
Biogeographic realms. Green: Nearctic, red: Palearctic, blue: Afrotropic, brown: Indomalaya, orange: Australasia, purple: Neotropic. Source: Wikimedia Commons.

**Figure A8.**
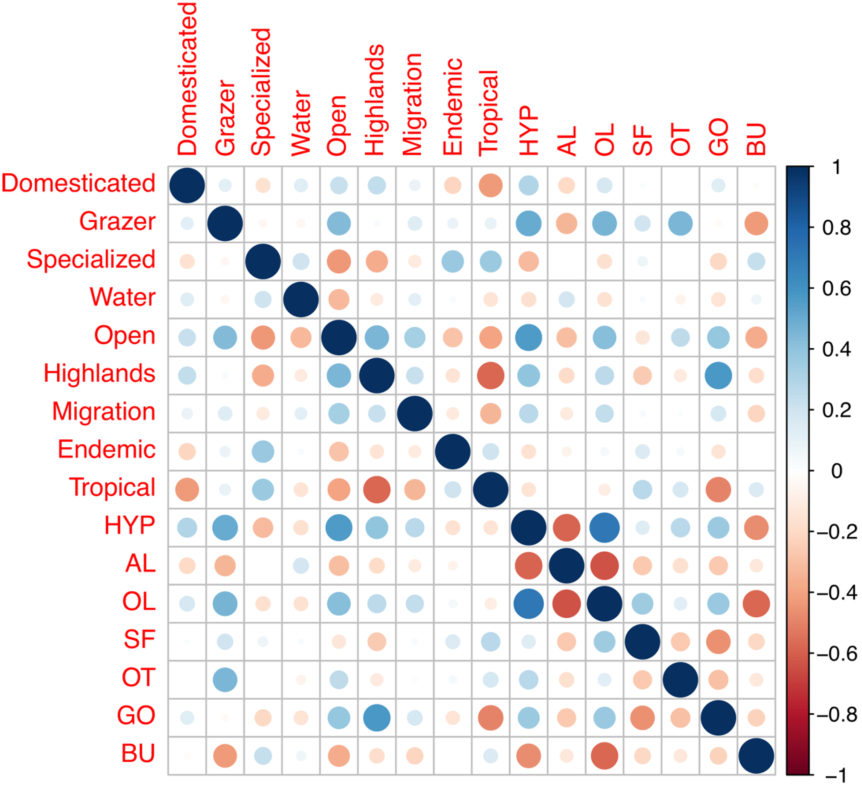
Correlations among the explanatory variables describing candidates for domestication (sample size is 68).

**Table A2.**
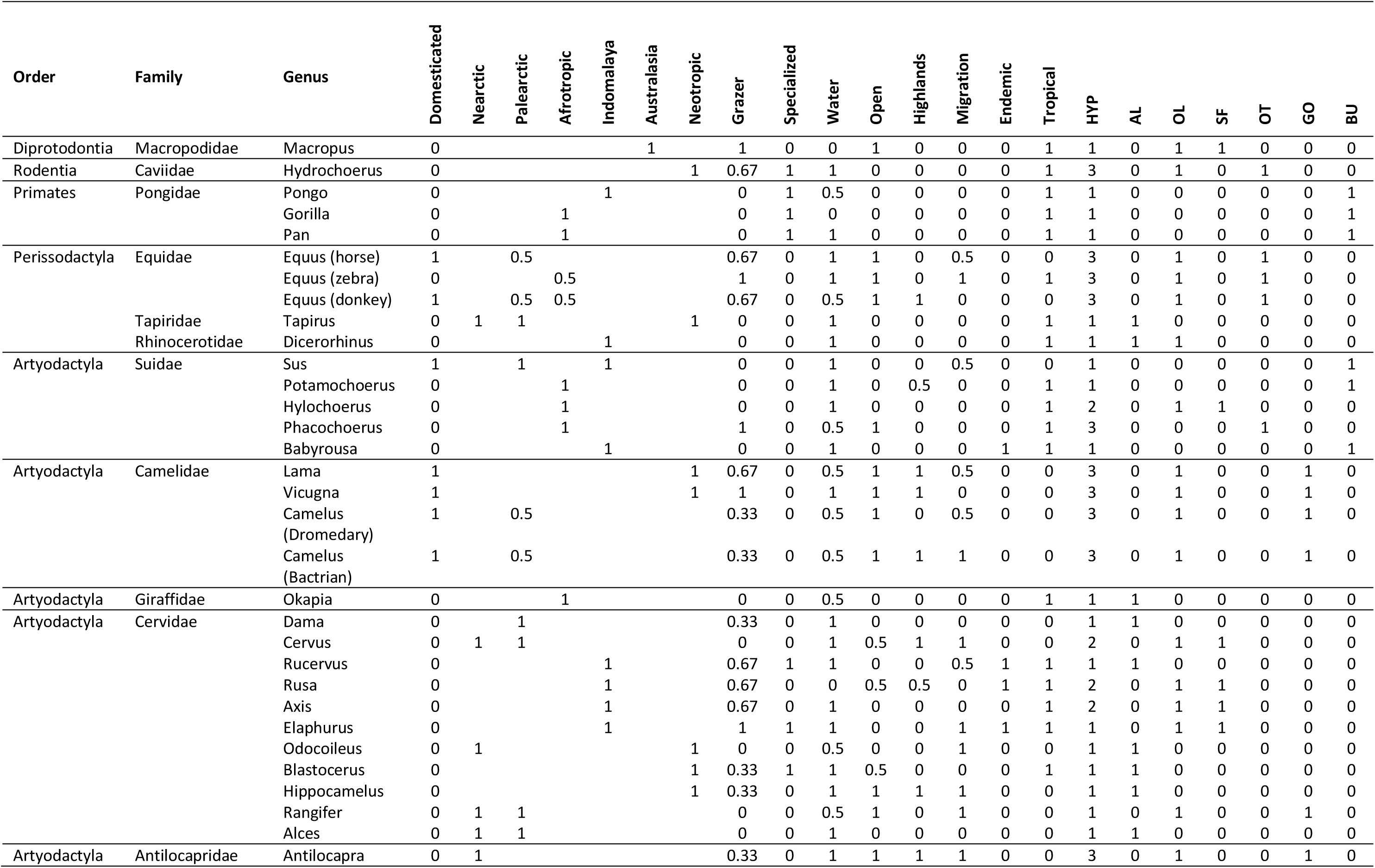
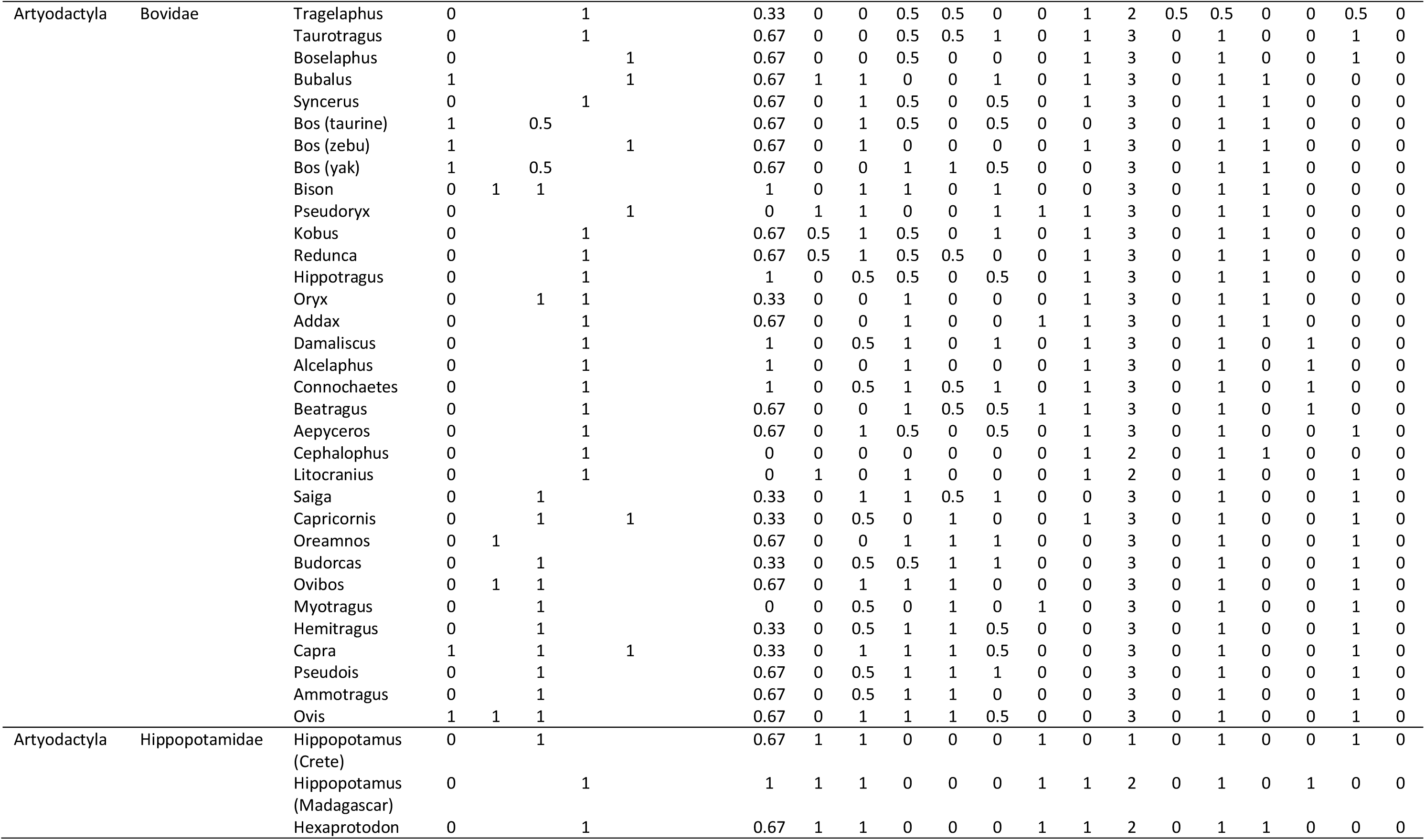
The dataset describing characteristics of candidate genera for domestication.

**Table A3.**
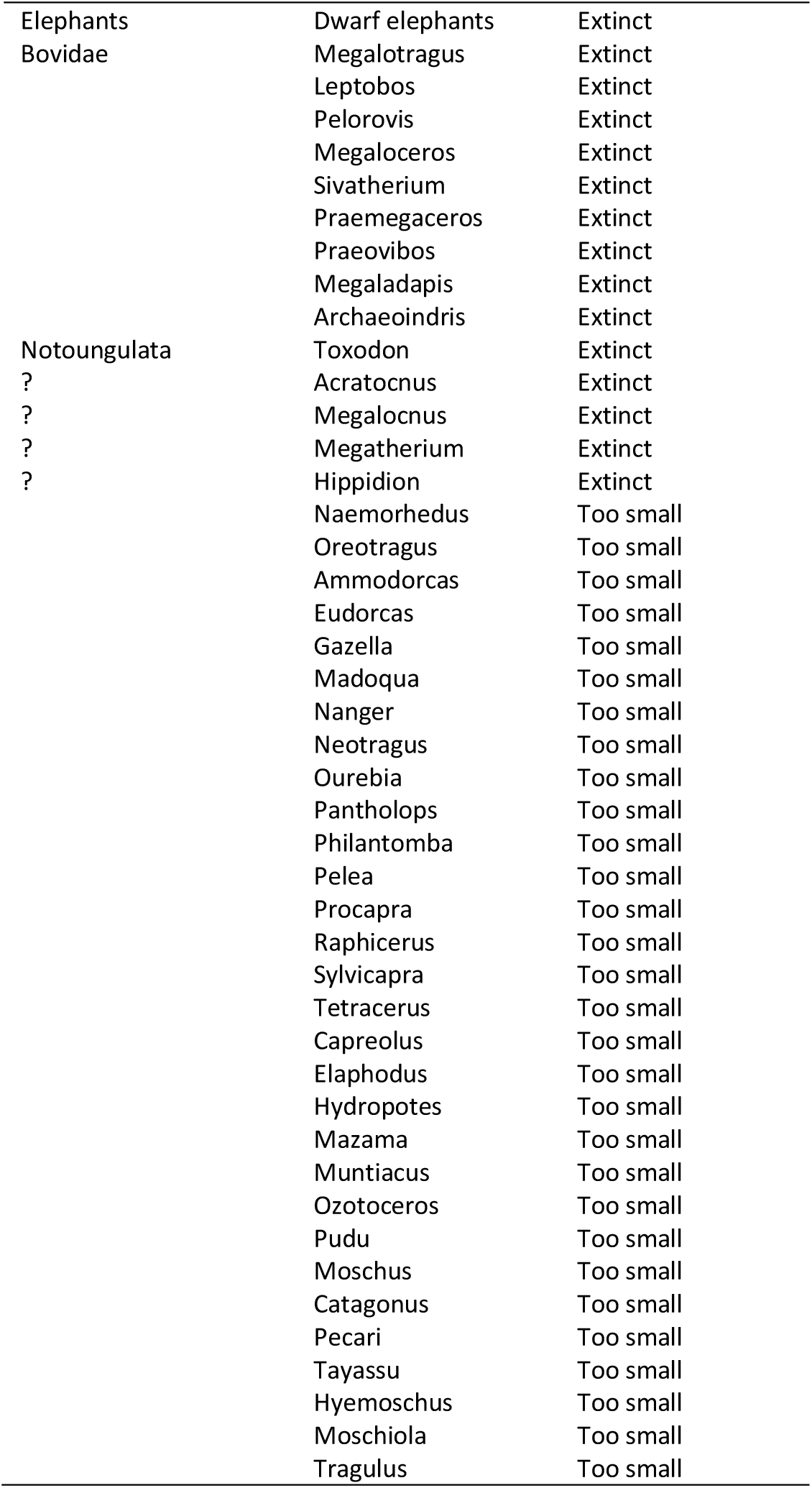
Genera excluded from the candidate list.

**Table A4.** First columns of the rotation matrix for PCA. Three largest weights within each component are highlighted.

| Variable | PC1 | PC2 | PC3 |
| --- | --- | --- | --- |
| Grazer | -0.245 | **0.365** | -0.204 |
| Specialized | 0.236 | 0.207 | 0.178 |
| Water | 0.125 | -0.045 | 0.195 |
| Open | **-0.388** | -0.055 | **-0.245** |
| Highlands | -0.299 | -0.308 | 0.107 |
| Migration | -0.205 | -0.063 | 0.122 |
| Endemic | 0.111 | 0.241 | 0.226 |
| Tropical | 0.221 | **0.405** | -0.126 |
| HYP | **-0.405** | 0.175 | 0.009 |
| AL | 0.251 | -0.247 | -0.197 |
| OL | **-0.368** | 0.263 | **0.257** |
| SF | 0.007 | **0.389** | 0.447 |
| OT | -0.092 | 0.260 | **-0.624** |
| GO | -0.285 | -0.316 | 0.199 |
| BU | 0.270 | -0.132 | -0.029 |

## Footnotes

1 The Fertile Crescent is a region in the Middle East, that lays across the territories of present-day Iraq, Israel, Syria, Lebanon, Egypt and Jordan, it reaches some parts of Turkey and Iran, and Cyprus may or may not be included. The name was popularized by James H. Breasted (1914).

2 https://github.com/zliobaite/teeth-domestication

3 A similar observation comes from isotope analysis of two grazing fossil pig genera that lived in Africa savannas: *Metridiochoerus*, which evolved locally, is thought to have been much less water-dependent than *Kolpochoerus* (Rannikko et al 2017), which originally came from Asia.

4 Hui Tang, personal communication, January 2018.

## REFERENCES

Bernor, R., Fahlbusch, V., Andrews, P., de Bruijn, H., Fortelius, M., Rogl, F., Steininger, F., Werdelin, L. (1996). The evolution of western Eurasian Neogene mammal faunas: a chronologic, systematic, biogeographic, and paleoenvironmental synthesis. In The Evolution of Western Eurasian Neogene Mammal Faunas, ed. R. Bernor, V. Fahlbusch, W. Mittmann, 449–70.

Bibi, F. (2013). A multi-calibrated mitochondrial phylogeny of extant Bovidae (Artiodactyla, Ruminantia) and the importance of the fossil record to systematics. BMC Evolutionary Biology 13 (1), 166.

Breasted, J.H. (1914). Earliest man, the Orient, Greece, and Rome. In Outlines of European history 1, ed. By J.H. Robinson, J.H Breasted and Ch.A. Beard, 56–57.

Crosby, A. (2006). Children of the sun: A History of humanitie’s unappeasable appetite for energy. Norton.

Childe, V.G. (1928). The Most Ancient Near East. London: Norton & Company.

Decory, M. (2019). A Universal Definition of ‘Domestication’ to Unleash Global Animal Welfare Progress. dA Derecho Animal: Forum of Animal Law Studies 10(2): 39–55.

Diamond, J. (1997). Guns, Germs, and Steel: The Fates of Human Societies. W. W. Norton & Company, New York, London.

Diamond, J. (2002). Evolution, consequences and future of plant and animal domestication. Nature 418, 700–707.

Eronen, J., Evans, A., Fortelius, M., Jernvall, J. (2011). Genera are often better than species for detecting evolutionary change in the fossil record: A reply to Salesa et al. Evolution, 65, 1514–1516.

Fortelius M, Eronen J, Jernvall J, Liu L, Pushkina D, Rinne J, Tesakov A, Vislobokova I, Zhang Z, and Zhou L. (2002). Fossil mammals resolve regional patterns of Eurasian climate change over 20 million years. Evolutionary Ecology Research 4: 1005–1016.

Fortelius, M., Getitz, S., Gyllenberg, M., Raia, P., Toivonen, J. (2015). Modeling the Population-Level Processes of Biodiversity Gain and Loss at Geological Timescales. American Naturalist 86(6):742–54.

Garrad, A. (1984). The selection of South-West Asian animal domesticates. Ed.: J. Clutton-Brock, C. Grigson. Animals and Archaeology 3 (S202), 117–132.

Guttal, V. and Couzin, I. D. (2010). Social interactions, information use, and the evolution of collective migration. PNAS 2010 107 (37), 16172–16177.

Hastie, T., Tibshirani, R., Friedman, J. (2009). The Elements of Statistical Learning: Data Mining, Inference, and Prediction. Second Edition. Springer.

Jarman, O. (1974). The social organisation of antelope in relation to their ecology. Behaviour, 48(3-4), 215–267.

Kottek, M., J. Grieser, C. Beck, B. Rudolf, and F. Rubel, 2006: World Map of the Köppen-Geiger climate classification updated. Meteorol. Z., 15, 259–263.

Larson, G., Piperno, D.R., Allaby, R.G., Purugganan, M.D., Andersson, L., Arroyo-Kalin, M., Barton, L. et al. (2014). Current perspectives and the future of domestication studies. PNAS 111 (17), 6139–6146.

Levin, S., Carpenter, H., Godfray, Ch., Kinzig, A., Loreau., M., Losos, J., Walker, B., Wilcove, D. (2012). Princeton guide to ecology. Princeton University Press.

Liow, L. H., Fortelius, M., Lintulaakso, K., Mannila, H., Stenseth, N. C. (2009). Lower extinction risk in sleep-or-hide mammals. The American Naturalist 173 (2), 264–272.

MacHugh, D.E., Larson, G., and Orlando, L. (2017). Taming the Past: Ancient DNA and the Study of Animal Domestication. Annu. Rev. Anim. Biosci. 5, 6.1–6.23.

Nowak, R. (2018). Walker’s Mammals of the World. Johns Hopkins University Press.

Oksanen, O., Zliobaite, I., Saarinen, J., Lawing, A. and Fortelius, M. (2019). A Humboldtian approach to life and climate of the geological past: estimating palaeotemperature from dental traits of mammalian communities. Submitted to the Journal of Biogeography 46(8): 1760–1776.

Olson, D. M., Dinerstein, E., Wikramanayake, E. D., Burgess, N. D., Powell, G. V. N., Underwood, E. C., D’Amico, J. A., Itoua, I., Strand, H. E., Morrison, J. C., Loucks, C. J., Allnutt, T. F., Ricketts, T. H., Kura, Y., Lamoreux, J. F., Wettengel, W. W., Hedao, P., Kassem, K. R. (2001). Terrestrial ecoregions of the world: a new map of life on Earth. Bioscience 51(11), 933–938.

Owen-Smith, N. and Mills, M. G. L. (2008). Predator–prey size relationships in an African large-mammal food web. Journal of Animal Ecology 77, 173–183.

Rabanus-Wallace, T., Wooller, M.J., Zazula, G.D., Shute, E., Jahren, A.H., Kosintsev, P., Burns, J.A., Breen, J., Llamas, B., and Cooper, A. (2017). Megafaunal isotopes reveal role of increased moisture on rangeland during Late Pleistocene extinctions. Nature Ecology & Evolution, 1, 0125.

Rannikko, J., Žliobaitė, I. and Fortelius, M. (2017). Relative abundances and palaeoecology of four suid genera in the Turkana Basin, Kenya, during late Miocene to Pleistocene. Palaeogeography, Palaeoclimatology, Palaeoecology 487, 187–193.

Ray, N. and Adams, J. (2001). A GIS-based vegetation map of the world at the Last Glacial Maximum (25,000-15,000 BP). Internet Archaeology 11.

The NOW Community (2018). New and Old Worlds Database of Fossil Mammals (NOW). Licensed under CC BY 4.0. Last retrieved on 1.10.2018 from http://www.helsinki.fi/science/now/.

Wilson and Reeder (2005). Mammal Species of the World: A Taxonomic and Geographic Reference. 3rd edition. Johns Hopkins University Press.

Witten, I., Frank, E., Hall, M., Pal, Ch. (2016). Data Mining: Practical Machine Learning Tools and Techniques. Elsevier.

Zliobaite, I., Fortelius, M., Stenseth, N. Chr. (2017). Reconciling taxon senescence with the Red Queen’s hypothesis. Nature 552, p. 92–95.

Zliobaite, I., Rinne, J., Toth, A., Mechenich, M., Liu, L., Behrensmeyer, A.K., Fortelius, M. (2016). Herbivore teeth predict climatic limits in Kenyan ecosystems. PNAS 113(45), p. 12751–12756.

Zeder, M. (2008). Domestication and early agriculture in the Mediterranean Basin: Origins, diffusion, and impact. PNAS 105 (33), 11597–11604.

Zeder, M.A. (2017). Domestication as a model system for the extended evolutionary synthesis. Interface Focus 7, 20160133.

## Supplementary references

Galbrun, E., Tang, H, Fortelius, M., Žliobaitė, I. (2018). Computational biomes: The ecometrics of large mammal teeth. Paleontologia Electronica, Article number: 21.1.3A.

Noe-Nygaard, N., Price, T.D., Hede, S.U. (2005). Diet of aurochs and early cattle in southern Scandinavia: evidence from ^15^N and ^13^C stable isotopes. Journal of Archaeological Science 32, 855–871.

Oksanen, O., Zliobaite, I., Saarinen, J., Lawing, A. and Fortelius, M. (2018). A Humboldtian approach to life and climate of the geological past: estimating palaeotemperature from dental traits of mammalian communities. Journal of Biogeography 46 (8), 1760–1776.

van Vuure, C. (2005). Retracing the Aurochs – History, Morphology and Ecology of an extinct wild Ox. Pensoft Publishers.

Žliobaitė, I., Tang, H., Saarinen, J., Fortelius, M., Rinne, J., Rannikko, J. (2018). Dental ecometrics of tropical Africa: linking vegetation types and communities of large plant-eating mammals. Evolutionary Ecology Research 19, p. 127147.

